# Comparative Evaluation of Solid-Phase and Membrane Mimetic Strategies in Membrane Proteome Coverage and Disease-State Analysis

**DOI:** 10.1101/2025.08.25.672181

**Authors:** Frank Antony, Ashim Bhattacharya, Hiroyuki Aoki, Rupinder S. Jandu, Abdualrahman M. Abdualkader, Rami Al Batran, Mohan Babu, Franck Duong van Hoa

**Author notes:** Equal contribution to the work (Frank Antony, Ashim Bhattacharya).

## Abstract

Membrane proteins (MPs) are vital to cellular signaling, metabolism, and disease pathology, yet remain underrepresented in proteomics. To address this, several independent workflows have been developed to enable the profiling of the membrane proteome, however the relative advantages and limitations of each method remain poorly defined. Here, we systematically compare four classical solid-phase membrane proteomic workflows (SP3, SP4, FASP, S-Trap) and three membrane mimetic strategies (Peptidisc, nanodisc, and SMALP copolymer) for mass spectrometry-based membrane proteome profiling, using healthy (LFD) and obese (HFD) mouse liver tissue. We found that the solid-phase methods yield higher total protein identifications, while the membrane mimetic systems enrich MPs. SMALP copolymer displays intermediate characteristics between the solid-phase and membrane mimetic workflows. Peptidisc and nanodisc stand out for their enrichment of MPs, although Peptidisc shows better enrichment of plasma membrane integral MPs, particularly those with 11+ transmembrane segments. In the context of HFD-induced liver proteome remodeling, the Peptidisc workflow outperformed the other six methods by capturing the highest number of differentially expressed MPs and demonstrating the greatest accuracy in detecting MP-level dysregulation. Collectively, this comparative analysis highlights the trade-offs between depth of proteome coverage and MP enrichment across workflows, underscoring the importance of method selection based on total protein counts, MP enrichment, and the accurate detection of MP-level dysregulation.

**Highlights:**

- Systematic comparison of seven workflows for membrane proteomics
- Solid-phase methods enrich soluble proteins; mimetics enrich membrane proteins
- SMALP displays intermediate performance between other workflows
- Peptidisc captures the most dysregulated membrane proteins in diseased liver
- Peptidisc most accurate in detecting membrane protein dysregulation

**In Brief Statement:** This study presents a systematic comparison of seven proteomic workflows for membrane protein profiling. Solid-phase methods yield higher total protein identifications, whereas membrane mimetics enrich more membrane proteins. Among tested methods on the diseased mouse liver, Peptidisc captures more differentially expressed membrane proteins and demonstrates superior accuracy in detecting membrane protein-level dysregulation. These findings provide a practical framework for selecting proteomic strategies tailored to membrane protein enrichment and biological insight.

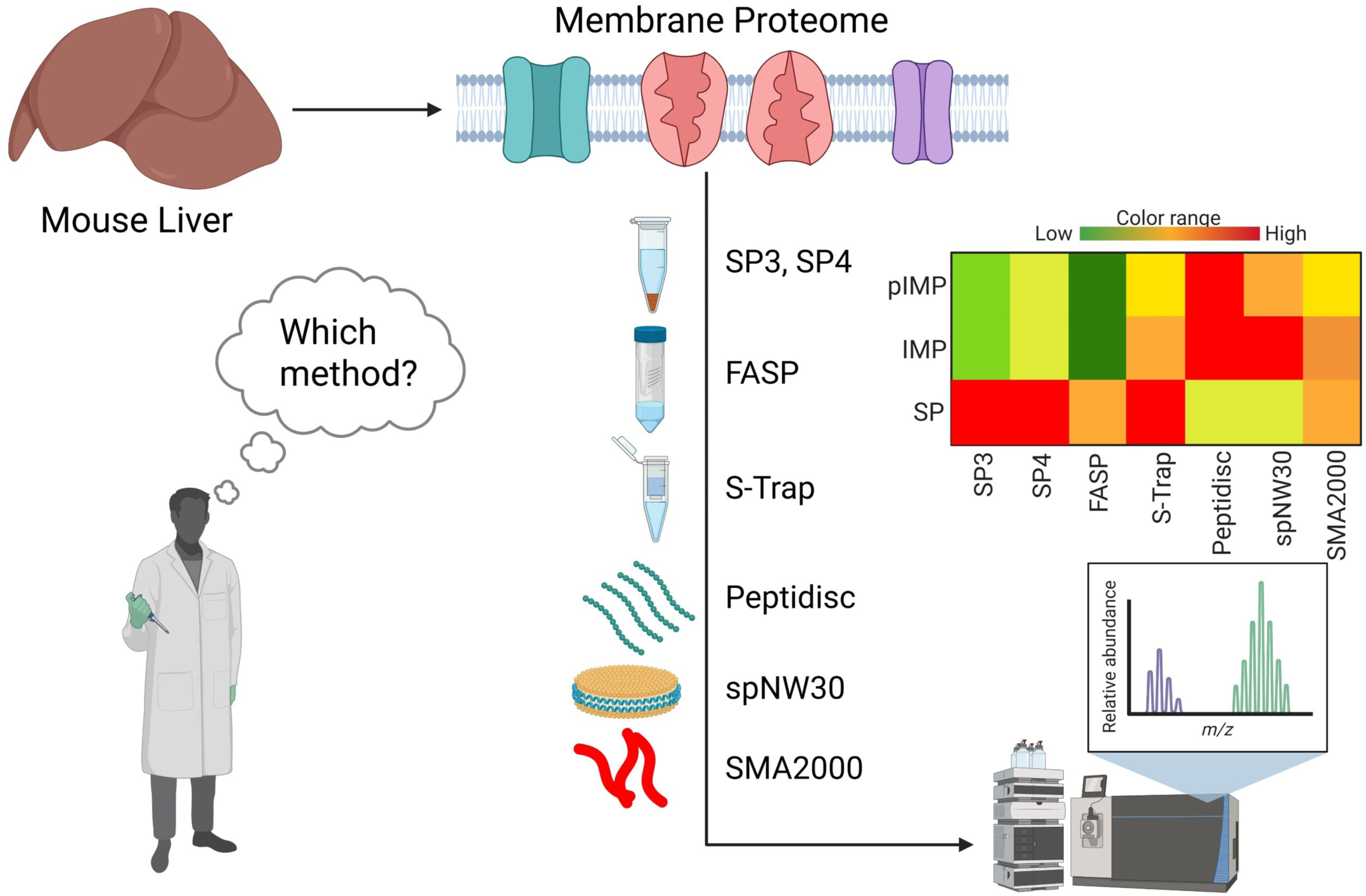

## Introduction

Membrane proteins (MPs) are central to cellular signaling, transport, and metabolic regulation, orchestrating essential biological processes such as receptor-mediated signaling, nutrient uptake, ion exchange, and detoxification(1). Their abundance and activity are critical for maintaining cell and tissue function(2), and their dysregulation has been implicated in a wide range of diseases, including neurodegenerative disorders(3,4), hypertension(5), cancer(6), muscular dystrophy(7), alcoholic fatty liver disease(8), and non-alcoholic fatty liver disease(9). Despite constituting over 30% of the human proteome(10) and more than ∼60% of current drug targets(11), MPs remain underrepresented in global proteomics datasets(12–14). Accurate identification and characterization of MPs is therefore essential for elucidating disease mechanisms, discovering biomarkers, and advancing drug development.

A major challenge in membrane proteomics lies in the efficient extraction of MPs from the lipid bilayer, a critical first step in bottom-up mass spectrometry (MS), which typically involves cell lysis, protein solubilization, denaturation, reduction, alkylation, enzymatic digestion, and peptide separation via liquid chromatography–tandem MS (LC–MS/MS)(15,16). Within this multi-step workflow, sample preparation is the critical step that governs the efficiency of protein capture and downstream analysis. This is pronounced for MPs, which are typically low in abundance, structurally complex, and prone to aggregation due to their hydrophobic transmembrane domains(17). To extract MPs, amphipathic detergents are commonly employed to disrupt lipid–protein interactions and solubilize membrane components. Strong ionic detergents such as SDS and RIPA are highly effective for membrane solubilization but are incompatible with downstream workflows, as they inhibit protease activity(18), compromise chromatographic separation(19), and suppress peptide ionization(20). Milder detergents like n-dodecyl-β-D-maltoside (DDM) and sodium deoxycholate (SDC) offer greater compatibility with proteolytic digestion and MS but are less effective at solubilizing MPs(21,22).

Critically, detergent removal prior to MS is essential but often leads to MP aggregation and precipitation(23). Most differences between membrane proteomic workflows stem from how detergents are removed and how proteins are stabilized post-extraction, which directly affects MP retention, digestion efficiency, and MS detection. As such, variability at this step is a major contributor to differences in membrane proteome coverage across sample preparation strategies.

A number of sample preparation strategies have been adapted to address the challenges of membrane proteome analysis. Among them, four solid-phase methods have stood out, unified by their use of a solid bead or column matrix to facilitate detergent removal. These include single-pot solid-phase-enhanced sample preparation (SP3)(24), solvent precipitation SP3 (SP4)(25), filter-aided sample preparation (FASP)(26), and suspension trapping (S-Trap)(27). SP3 and SP4 both involve protein aggregation onto beads to remove detergents. The distinction lies in the bead type: SP3 uses carboxylated magnetic beads, whereas SP4 employs silica glass beads.

FASP utilizes a 10-30 kDa ultrafiltration filter to capture SDS-denatured proteins while washing away detergent micelles. Meanwhile, S-Trap, proteins are acidified with phosphoric acid and captured on a quartz fiber matrix using a methanol-triethylammonium bicarbonate (TEAB) solution followed by washes to remove contaminants.

Importantly, while these techniques effectively remove detergents, their underlying mechanisms, such as organic solvent-induced precipitation (SP3 and SP4) or filtration-based retention (FASP and S-Trap), favor the capture and digestion of soluble, hydrophilic proteins. MPs, by contrast, are prone to aggregation under these conditions and are often retained in forms less accessible to proteases, resulting in incomplete digestion and poor MS detection(15).

Additionally, in these workflows, peptides are generated directly from crude membrane extracts without a dedicated MP enrichment step. Consequently, solid-phase methods remain biased toward soluble proteins, limiting their ability to comprehensively profile the membrane proteome and its roles in health and disease.

Despite advances in solid-phase methodology, MPs remain challenging targets for proteomic analysis, and more broadly, for biochemistry and structural biology(15). To facilitate their capture and study, researchers have developed membrane mimetics, amphipathic scaffolds that stabilize MPs by wrapping around their hydrophobic transmembrane regions, rendering them fully water-soluble(28). By preserving MPs in a native state, membrane mimetics minimize aggregation circumventing issues observed with solid-phase methods(15). In addition to enhancing MP stability, they can also facilitate the removal of soluble protein contaminants through affinity-tagging of the scaffold, enabling an additional MP enrichment step prior to MS analysis.

Among them, nanodiscs, composed of discoidal lipid bilayers encircled by a belt of two apolipoprotein derived membrane scaffold proteins (MSP), offer a robust approach for reconstituting MPs into soluble assemblies(29). Additionally, the MSP can be His-tagged, enabling affinity enrichment of the MSP-captured membrane proteome(30). A promising advancement is the 30 nm diameter SpyTag/SpyCatcher circularized nanodisc (spNW30)(31). Unlike conventional MSP nanodiscs, which exhibit size heterogeneity at larger diameters (>16 nm), spNW30 achieves improved reconstitution yield, uniformity and stability by covalently linking the MSP’s N- and C-termini to form a closed ring. Additionally, its larger size enables the study of MP complexes and lipid-mediated processes requiring extended bilayer surfaces(31). However, the performance of nanodiscs for comprehensive membrane proteome profiling has yet to be evaluated.

Another previously adopted membrane mimetic platform for membrane proteome profiling is the styrene–maleic acid copolymer (SMA). This reagent spontaneously inserts into biological membranes and extracts MPs along with surrounding lipids, forming nanodiscs approximately ∼8-15 nm in diameter, termed SMA lipoparticles (SMALPs)(32). Unlike other membrane mimetics, SMALPs production requires no detergent at any stage of the workflow(33). In a benchmark study, Brown et al. evaluated several SMALP formulations across diverse tissue types and found SMA2000 (2:1 styrene-to-maleic acid copolymer) to robustly capture the membrane proteome(34). However, its performance relative to other membrane-mimetic systems, has yet to be evaluated.

Finally, the Peptidisc, composed of short amphipathic peptides, has emerged as another versatile membrane mimetic capable of reconstituting a broad range of detergent-solubilized MPs into stable, water-soluble particles(35). The addition of affinity tags, such as His or biotin, to the peptide scaffold has proved useful in enriching MPs prior to MS analysis(36,37). We recently demonstrated the robustness of this approach for membrane proteome profiling by capturing MPs from mouse brain, heart, kidney, lung, and liver(38). In a subsequent study, we showed that Peptidisc-based membrane proteomics enables identification of proteome dysregulation between normal and high-fat + ethanol-fed mouse livers (mLiver)(8). However, the Peptidisc has yet to be compared to other membrane mimetic systems in the context of membrane proteome profiling.

Despite the diverse repertoire of methods available for membrane proteome analyses, a systematic, side-by-side comparison of their workflows and performance has yet to be conducted. To address this gap, we systematically compared seven methods based on their ability to capture and enrich MPs from the mLiver. To further assess each method’s performance in detecting MP-level dysregulation under pathological conditions, we applied them to an obesogenic-diet-induced mouse model of metabolic dysfunction–associated steatotic liver disease (MASLD). MASLD, which arises from excessive lipid accumulation in hepatocytes, is the most common chronic liver disease globally(39). Given the liver’s reliance on MPs for nutrient sensing, lipid trafficking, and redox homeostasis(37,38), MP dysregulation is expected to be well detected in this diet-induced mLiver disease, making it a compelling model for benchmarking membrane proteomics workflows.

Herein, we present the first systematic comparison of four solid-phase (SP3, SP4, S-Trap, FASP) and three membrane mimetic (Peptidisc, spNW30, SMA2000) workflows, applied to both healthy and diseased mLiver tissue. We evaluated each method’s capacity to capture and enrich MPs, as well as to detect proteome-level dysregulation. By clarifying the trade-offs across workflows, our study aims to facilitate broader exploration of membrane-centric biology.

### Experimental Procedures

#### Materials

DDM and SDS were purchased from Anatrace. His6-tagged Peptidisc (purity >90%) were obtained from Peptidisc Biotech. Nickel nitrilotriacetic acid (Ni-NTA) chelating Sepharose resin was obtained from Qiagen. Silica beads and the complete protease inhibitor cocktail were purchased from Sigma. Carboxylate-modified paramagnetic beads (Sera-Mag Speed-Beads (Hydrophilic), CAT# 45152105050250, and Sera-Mag SpeedBeads (Hydrophobic), CAT# 65152105050250) were purchased from GE Healthcare (Chicago, IL). Amicon ultra centrifugal filters 30KDa and 100KDa molecular weight cut-off (MWCO) were from Millipore. S-Trap microcolumns were purchased from Protifi, USA. TEAB, phosphoric acid (HPLC grade), acetonitrile (99.9%), MS-grade trypsin and DNAase was purchased from Thermo Fisher Scientific. Octadecyl C18 Empore disks were purchased from 3M and Polygoprep 300-20 C18 powder was purchased from Macherey-Nagel. All other general reagents such as NaCl, Tris-base, DTT, PMSF, iodoacetamide (IAA), urea, organic solvents and acids were obtained from Bioshop or Fisher Scientific Canada.

#### Mouse treatment and harvest of organs

Animal experiments were approved by the Animal Experimentation Ethics Committees of the Université de Montréal (approval number 20-062) and performed in accordance with the Guide for the Canadian Council on Animal Care. Animals were housed in a 22^◦^C temperature-controlled unit under a 12 h–12 h light–dark cycle with standard environmental enrichment and *ad libitum* access to drinking water and food. Seven-week-old male C57BL/6J mice (The Jackson Laboratory) were fed either a low-fat diet (LFD) (10% kcal from lard; Research Diets D12450J) and used as the lean control group for comparison or a high-fat diet (HFD) (60% kcal from lard; Research Diets D12492) for 18 weeks to induce obesity and hepatic steatosis. The LFD contained 10% of its calories from fat and 70% from carbohydrates. The HFD contained 60% of its calories from fat and 20% from carbohydrates. The macronutrient sources remained the same for all diets, including the protein source (casein and L-cystine) and the carbohydrate source (corn starch, maltodextrin 10 and sucrose). Exactly 12 weeks post dietary implementation, mice were killed during the fed state using an I.P. injection of sodium pentobarbital (12 mg), after which tissues were extracted, rinsed five times in ice-cold PBS to remove blood and immediately snap frozen in liquid nitrogen stored at −80°C until processed.

#### Tissue processing and preparation of the membrane fraction

Frozen tissue samples were thawed on ice, minced, and subsequently homogenized in hypotonic lysis buffer (10 mM Tris-HCl pH 7.4; 30 mM NaCl, and 1 mM EDTA, 1x cocktail protease inhibitor and 1 mM PMSF). The homogenate was further incubated for 10 min on ice in the presence of 10 mM MgCl2 and 50 µg/ml DNAse. The enlarged cell suspension was lysed using a French press with 3 passages at 500 pounds per square inch (psi). The cell lysate was centrifuged at 1,200 x g for 10 min at 4^◦^C to remove unbroken cells and nucleus fraction. The supernatant was collected and centrifuged at 5,000 x g for 10 min at 4^◦^C to remove the mitochondrial fraction and the supernatant was ultracentrifuged at 110,000 x g for 45 min at 4^◦^C in a Beckman TLA110 rotor. The pellet was resuspended in 200 μL of TSG buffer (50 mM Tris, pH 7.9, 100 mM NaCl, 10% glycerol) to obtain a membrane fraction called “crude membrane” and stored at −80°C until use.

#### Sample preparation for SP3-based analysis

Sample preparation using the SP3 method was performed as previously described(24) with minor modifications. Briefly, a 1:1 ratio mixture of Sera-Mag SpeedBeads was washed three times with water using a magnetic rack and resuspended in water at a final concentration of 50 mg/mL. Crude membrane fractions (∼1 mg) were solubilized in ice-cold TS buffer (50 mM Tris-HCl, pH 8.0, 100 mM NaCl) supplemented with 1% (w/v) SDS for 30 min at 4°C with gentle shaking.

The solubilized proteins were clarified by ultracentrifugation at 110,000 × g for 15 min at 4°C. Prepared beads were added to 100 µg protein sample in a ratio of 1:10 (μg of protein/μg of SP3 beads). To induce binding of protein to beads, chilled acetonitrile (4°C) was added in a 4:1 (v/v) acetonitrile-to-protein ratio and were incubated at 24°C for 5 min with mixing at 1,000 rpm in a ThermoMixer. Tubes were placed in a magnetic rack and incubated for 5 min at RT. The supernatant was discarded and the beads were then rinsed three times with 180 µL of 80% ethanol, with pipette mixing between each wash. After the final wash, beads were briefly air-dried before proceeding with protein denaturation using 8M urea. The samples were sonicated in a water bath for 1min to fully disaggregate the beads. Further, disulfide bond reduction using 10 mM DTT for 1hr and alkylation with 20 mM IAA in the dark for 30 mins, followed by quenching with an additional 10 mM DTT for 30 mins and finally the urea was diluted to 1M with 50 mM ammonium bicarbonate, pH 8.0. Proteins were digested with trypsin at a 1:100 enzyme-to-protein ratio and incubated on shaker for 24 h at 25°C. Following digestion, samples were centrifuged at 20,000 × g for 1 min at 24°C. The supernatant was recovered by magnetic separation of the beads and transferred to fresh tubes. Peptides were acidified, desalted using C18 StageTips and were eluted with 40% acetonitrile containing 0.1% formic acid. Eluted samples were dried by vacuum centrifugation for subsequent Liquid chromatography-tandem mass spectrometry (LC-MS/MS) analysis.

#### Sample preparation for SP4-based analysis

The SP4 protocol was performed as previously described(25), with minor modifications. Briefly, silica beads (9–13 μm diameter) were resuspended in water, washed once with 100% acetonitrile, rinsed twice with water and resuspended at 50 mg/ml final. After each wash, the beads were isolated by brief centrifugation at 16,000 x g for 1 min. Crude membranes (∼1 mg) were resuspended in ice-cold TS buffer supplemented with 1% (w/v) SDS for 30 min at 4^◦^C with gentle shaking. The detergent extract was clarified by ultracentrifugation (110,000 x g for 15 min at 4^◦^C) and aliquots (100 µg each) were gently vortex-mixed with glass beads (1 mg).

Acetonitrile was then added to a final concentration of 80% and samples were centrifuged at 16,000 x g for 5 min. The beads were rinsed three times with 500 µl of 80% ethanol without disturbing the pellet. After a final wash, the beads were resuspended in 100 µl of 8 M urea and placed in a sonicator bath for 5 min. Proteins were reduced with 10 mM DTT for 1 hour, alkylated in the dark with 20 mM iodoacetamide and further reduced with 10 mM DTT for 30 min. The urea was diluted to 1 M with the 50mM ammonium bicarbonate buffer pH 8 and trypsin was added at an enzyme/protein ratio of 1:100 for 24 h at 25°C on a shaker. The beads were recovered by centrifugation at 16,000 x g for 1 min and the supernatant (800 µL) was placed in a new tube. The samples were acidified with 10% formic acid to pH 3 (35 µL) and loaded onto hand-packed C18 Stage-tips. Samples were eluted with 40% Acetonitrile containing 0.1% formic acid. The eluted peptides were dried by vacuum centrifugation.

#### Sample preparation for FASP-based analysis

FASP was performed using a modified version of a previously described protocol(40). Briefly, crude membrane fractions (∼1 mg) were solubilized in TS buffer containing 1% SDS and 100 mM dithiothreitol (DTT) for 30 min at 4°C. Insoluble material was removed by ultracentrifugation at 55,000 rpm for 15 min at 4°C (TLA 55 rotor), and the concentration of solubilized proteins was determined. Aliquots containing 100 µg of protein were mixed with 200 μL of 8 M urea in 100 mM ammonium bicarbonate pH 8.0 (8M urea-NH4HCO3) and transferred to Amicon Ultra centrifugal filter units (30 kDa MWCO) and centrifuged at 14,000 × g for 15 min at 20°C. The eluates were discarded, and samples were washed twice 100 μL of 8M urea-NH4HCO3 and the units were centrifuged again. Proteins were then alkylated by adding 100 µL of 50 mM iodoacetamide in 8M urea-NH4HCO3, incubated in the dark at room temperature for 30 min, and centrifuged again. Subsequently, the filters were washed once with 100 µL of 8M urea-NH4HCO3, followed by two washes with 100 µL of 20 mM ammonium bicarbonate.

Proteins were digested for 24 h at 25°C with trypsin at an enzyme-to-protein ratio of 1:25 in 80 µL of 20 mM ammonium bicarbonate. Digested peptides were collected by centrifugation at 14,000 × g for 10 min, followed by an additional rinse with 100 µL of 20 mM ammonium bicarbonate. Flow-throughs were pooled. Further, the samples were acidified to pH ∼3 using 10% formic acid. Peptides were desalted using C18 stagetips. Peptides were eluted with 40% acetonitrile containing 0.1% formic acid and dried by vacuum centrifugation (SpeedVac, no heat) prior to LC-MS/MS analysis.

#### Sample preparation for S-Trap based analysis

Approximately 1 mg of crude membrane fractions were solubilized in ice-cold TS buffer containing 1% (w/v) SDS. The mixture was gently shaken at 4 °C for 30 minutes, then subjected to ultracentrifugation at 110,000 × g for 15 minutes at 4 °C to clarify the solubilized proteins.

Protein digestion in the S-Trap filter was performed following the manufacturer’s procedure. Briefly, 100µg of total protein was reduced with 10 mM DTT for 1 hour, alkylated with 20 mM IAA in the dark for 30 min and reduced with 10 mM DTT for 30 min. The sample was added with a final concentration of 1.2% phosphoric acid and then six volumes of binding buffer (90% methanol; 100 mM TEAB; pH 7.1). After gentle mixing, the protein solution was loaded to an S-Trap filter, spun at 10,000 x g for 30 sec. Filters were washed three times with 150μL of binding buffer and centrifuged at 10,000 g for 30 sec with 180° rotation of the filters between washes.

Samples were then digested with trypsin (1:50 trypsin/protein) for 24 h at 25°C. The digested peptides were eluted using 50 mM ammonium bicarbonate; trypsin was added to this fraction and incubated for 24 h at 25°C. Two more elutions were made using 0.2% formic acid and 0.2% formic acid in 50% acetonitrile. The three elutions were pooled together and were desalted using C18 stagetips. Peptides were eluted with 40% acetonitrile containing 0.1% formic acid and vacuum dried prior to LC-MS/MS analysis.

#### Sample preparation for Peptidisc-based analysis

Crude membranes (∼1 mg) were resuspended in ice-cold TS buffer supplemented with 1% (w/v) DDM for 30 min at 4^◦^C with gentle shaking. The detergent extract was clarified by ultracentrifugation (110,000 x g for 15 min at 4^◦^C) and 1 mg aliquots (500 µl) were incubated with a 3-fold excess of His6-tagged Peptidiscs for 15 min at 4^◦^C. The sample was rapidly diluted to 5 mL in the TS buffer placed in a 100 kDa cutoff centrifugal filter. The sample was concentrated at 3,000 x g, 10 minutes to approximately 200 µl and diluted again to 5 ml, followed by another concentration step to approximately 200 µl. The reconstituted start Peptidisc library (4 mg total in 1 mL) was incubated with 60 μL of Ni-NTA chelating Sepharose for 1 hour at 4^◦^C with shaking. Following five thorough washes with TS buffer 1 mL each, the Peptidisc library was eluted in 150 μL of TS buffer containing 600 mM imidazole, recovering 1 mg at a concentration of ∼6.5 µg/µL. All fractions from the reconstitution through the purification steps were subjected to analysis by 15% SDS PAGE, followed by Coomassie blue staining of the gel. An aliquot of the purified MP library containing (∼100 μg; ∼16 µL) was treated with 6 M urea at room temperature for 30 min, followed by reduction with 10 mM DTT for 1 hour. Alkylation was performed with 20 mM IAA in the dark at room temperature for 30 min, followed by re-addition of 10 mM DTT for 30 min. The urea concentration was diluted to 1 M with 50 mM ammonium bicarbonate, pH 8.0 and trypsin was added at an enzyme/protein ratio of 1:100 for 24 h at 25°C on a shaker. Subsequently, the digested peptides were acidified to pH 3 with the addition of ∼20µl 10% formic acid and desalted using hand-packed Stage-Tips C18. The eluted peptides were dried by vacuum centrifugation.

#### Sample preparation for spNW30-based analysis

Approximately 1 mg crude membranes were solubilized in 1% DDM for 30 min at 4°C with gentle shaking. The insoluble material was pelleted by ultracentrifugation (110,000 x g, 15 min). The detergent extract (500 μL) was then reconstituted by mixing it with a 1:1 ratio of His-spNW30 for 15 min at 4°C. The sample was rapidly diluted to 5 mL in TS buffer over a 100 kDa MWCO centrifugal filter. The sample was concentrated (3000 x g, 10 min) to ∼200 μL and diluted again to 5 mL. After another round of concentration to ∼200 μL, the spNW30 start library was incubated with 60 μL of Ni-NTA resin for 1 h at 4 °C with shaking. After extensive washing with TS buffer (5 washes, 1 mL each), the “Purified Library” was eluted in 150 μL of TS buffer supplemented with 600 mM imidazole. Protein aliquots of all fractions from the reconstitution to the purification stages were analyzed by 15% SDS PAGE, followed by Coomassie blue staining of the gel.

#### Sample preparation for SMA2000-based analysis

The SMA polymer with a 2:1 styrene to maleic acid ratio was prepared according to a previously described protocol(41). Briefly, a 10% solution of SMA 2000 (Cray Valley) was refluxed in 1 M KOH at 80°C for 3 hours, leading to full solubilization of the polymer. The polymer was subsequently precipitated by gradually adding 6 M HCl with continuous stirring, followed by centrifugation at 1500 × g for 5 minutes to collect the pellet. The pellet was washed three times with 50 mL of 25 mM HCl, followed by an additional wash with ultrapure water, and then lyophilize. The SMA polymer pellet was subsequently re-dissolved at a concentration of 10% w/v in 25 mM Tris-HCl, with a final the pH adjusted to 8.0. 100 µg crude membrane fractions were solubilized in TS buffer containing 2.5% SMA2000 for 4hrs at 4°C, clarified by ultracentrifugation at100,000 x g for 15 min at 4°C. The solubilized samples were methyl tert-butyl ether (MTBE) extracted as previously described(34). In brief, 200 µl protein sample was mixed with 1ml ice cold MTBE:Methanol (3:1) and vortexed for 45 min at 4 °C. The samples were sonicated in Branson bath-sonicator for 15 minutes at 4 °C, followed by the addition of 650 μl of a 3:1 H₂O:Methanol mixture containing 1% KCl. The precipitated proteins were pelleted by centrifugation at 20,000g for 5 min. The supernatant was discarded, and the protein pellet was washed twice with 80% Methanol followed by centrifugation at 20,000g for 5 min. The supernatant was aspirated away and the pellet was dried under vacuum. The pellet was resuspended in 100 µl of 8 M urea and placed in a sonicator bath for 5 min. Proteins were reduced with 10mM DTT for 1 hour, alkylated in the dark with 20mM IAA in the dark for 30 mins, followed by quenching with an additional 10 mM DTT for 30 mins. The urea was diluted to 1 M with the 50mM ammonium bicarbonate buffer pH 8 and trypsin was added at a 1:100 enzyme-to-protein ratio for digestion over 24 hours at 25°C with continuous shaking. Following digestion, peptides were acidified, desalted using C18 StageTips and then eluted with 40% Acetonitrile containing 0.1% formic acid. The eluted samples were subsequently dried using vacuum centrifugation for subsequent LC-MS/MS analysis.

#### LC and MS/MS analyses

NanoLC connected to an Orbitrap Exploris mass spectrometer (Thermo Fisher Scientific) was used for the analysis of all samples. The peptide separation was carried out using a Proxeon EASY nLC 1200 System (Thermo Fisher Scientific) fitted with a custom-made C18 column (15 cm x 150 μm ID) packed with HxSil C18 3 μm Resin 100 Å (Hamilton). A gradient of water/acetonitrile/0.1% formic acid was employed for chromatography. The samples were injected onto the column and run for 180 minutes at a flow rate of 0.60 μl/min. The peptide separation began with 1% acetonitrile, increasing to 3% in the first 4 minutes, followed by a linear gradient from 3% to 23% acetonitrile over 86 minutes, then another increase from 24% to 80% acetonitrile over 35 minutes, and finally a 35-minute wash at 80% acetonitrile, and then decreasing to 1% acetonitrile for 10 min and kept 1% acetonitrile for another 10 min. The eluted peptides were ionized using positive nanoelectrospray ionization (NSI) and directly introduced into the mass spectrometer with an ion source temperature set at 250°C and an ion spray voltage of 2.1 kV. Full-scan MS spectra (m/z 350–2000) were captured in Orbitrap Exploris at a resolution of 120,000 (m/z 400). The automatic gain control was set to 1e6 for full FTMS scans and 5e4 for MS/MS scans. Ions with intensities above 1500 counts underwent fragmentation via NSI in the linear ion trap. The top 15 most intense ions with charge states of ≥2 were sequentially isolated and fragmented using normalized collision energy of 30%, activation Q of 0.250, and an activation time of 10 ms. Ions selected for MS/MS were excluded from further selection for 3 seconds. The Orbitrap Exploris mass spectrometer was operated using Thermo XCalibur software.

#### Data Analysis in Fragpipe

The label free quantification (LFQ) of raw MS data was processed using the MSFragger (v 4.1) in FragPipe (v 22.0) (42,43) using LFQ-MBR workflow and identification search was performed against UniProt-mouse protein database (UP000000589, December 2024, 54,727 entries) concatenated with reverse decoy database. Precursor and fragment mass tolerance were both set to 20 ppm. Digestion settings ensured usage of the “stricttrypsin” enzyme, focusing on fully tryptic peptides (with up to 2 missed cleavages), within a peptide length range of 7–50 and mass range of 500–5000. Carbamidomethylation of cysteines was set as fixed modification, and methionine oxidation and acetylation of the N terminus were set as variable modification. A maximum variable modification per peptide was set to three. For validation, MSBooster in combination with Percolator was enabled, and a false discovery rate (FDR) of 1% was applied at both the peptide spectrum match (PSM) and protein levels (sequential filtering), utilizing the corresponding decoy sequences for FDR estimation. At the quantification stage, enabled MaxLFQ intensity, match-between-runs aligning with FDR 1% at ion and normalize intensities across runs, adopting the peptide-protein uniqueness option as “unique plus razor”.

Quantification was performed after a 1% protein-level FDR filtering, using IonQuant with default settings (10 PPM precursor mass tolerance, 0.4 min retention time tolerance, 0.05 1/K0 ion mobility tolerance, top 3 ions) with both match-between-runs and MaxLFQ enabled.

#### Protein annotation

The protein list obtained from FragPipe was subjected to a gene ontology (GO)-term analysis using the UniProtKB database to identify proteins with the GO-term “membrane”. Within this group, proteins containing at least one α-helical transmembrane segment were labeled as IMPs. While those without any transmembrane segment were annotated as MAPs. The Phobius web server was utilized (http://phobius.sbc.su.se/) to predict the number of transmembrane segments(44). The subcellular localization of the IMPs was further classified using the GO-term “Subcellular location [CC].” These were divided into pIMPs if they contain the GO-term ‘plasma membrane’. Alternatively, they were labeled as oIMPs when containing keywords such as ER, Golgi membrane, vesicle membrane (including endosome, exosome, peroxisome, lysosome, and vesicle), mitochondrial and nucleus membranes).

#### Figure Generation, Data Analysis, and Code Availability

The graphical abstract and Figure 1 were generated with Biorender. Figures 2-5 and Supplemental Figures 2-4 6-7 were generated using R (v4.3.0). Statistical analysis for Supplemental Figure 4 was performed with R. Volcano plots statistical analysis p-value calculations for Figures 4-6 were performed using Perseus. Figures 6-7 and Supplemental Figure 5 were generated with GraphPad Prism 10. Figure 8 Gene Ontology term analysis was performed using the DAVID Bioinformatics Resources, and the results were visualized in Microsoft Excel. Custom scripts used for data processing and visualization are available from the corresponding author upon reasonable request.

**Figure 1:**
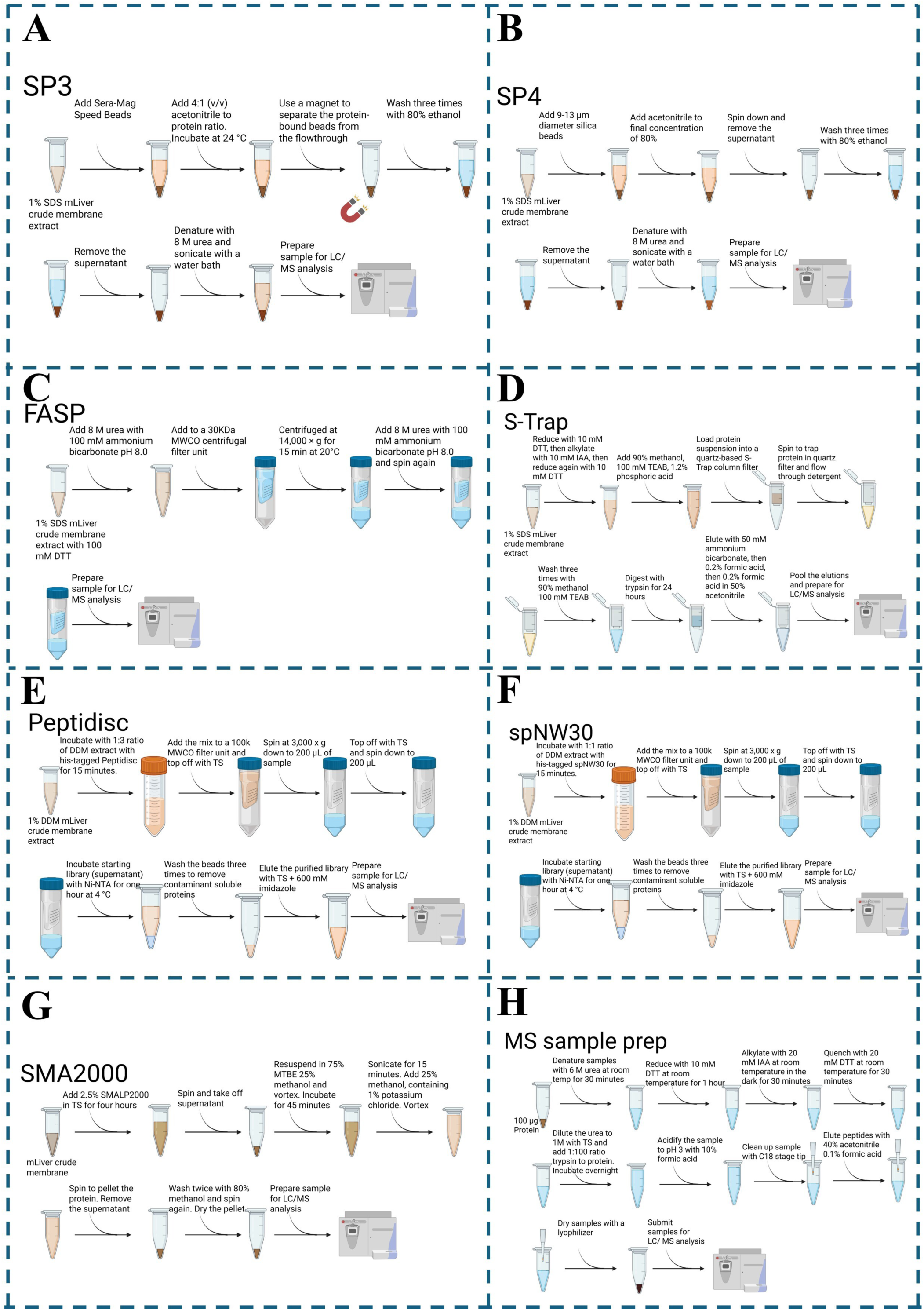
Methodological Schematic of the Seven Proteomic Workflow and the MS Sample Preparation Protocol. (A-H) Figure summarizing the (A) SP3, (B) SP4, (C) FASP, (D) S-TRAP, (E) Peptidisc, (F) spNW30, (G) SMA2000, (H) and MS sample preparation workflow as performed in this study. Figure was generated with Biorender.

**Figure 2.**
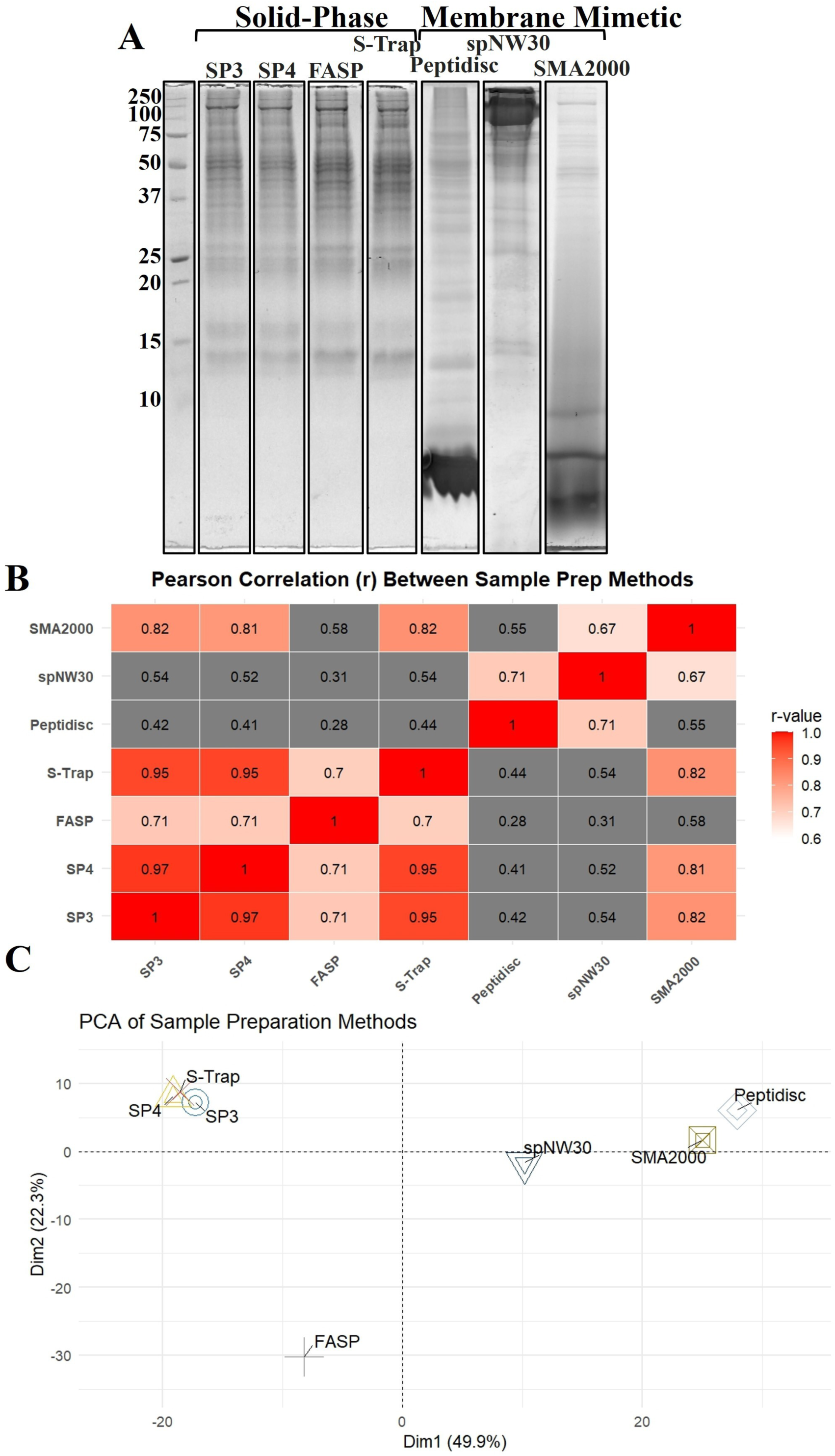
SDS-Page Analysis and Inter-Method Correlation Between the Workflows. (A) SDS-PAGE (15%) analysis of mLiver extracts prepared using the different workflows. A molecular weight marker is shown on the left. Gels were stained with Coomassie Blue and imaged using the IR-short channel of the Amersham Typhoon imager. **(B)** Correlation matrix showing pairwise Pearson correlation coefficients (r-values) between the sample preparation methods (SP3, SP4, FASP, S-Trap, Peptidisc, spNW30, SMA2000) based on MaxLFQ intensities from the replicate with the highest number of protein identifications per method. Correlation coefficients were calculated using the cor() function in R with method = “pearson” and use = “pairwise.complete.obs”. Visualization was performed using the ggplot2 and reshape2 packages, with color intensity indicating correlation strength. (C) PCA of the same MaxLFQ intensity dataset in (A), where each method is represented as a point based on its global protein intensity profile. Protein intensities were log₂-transformed, centered, and scaled before PCA using the prcomp() function. The PCA plot was generated with the factoextra and ggrepel packages, showing separation of sample preparation strategies in reduced dimensional space. Method names are labeled on the plot, and color-coded by workflow.

**Figure 3.**
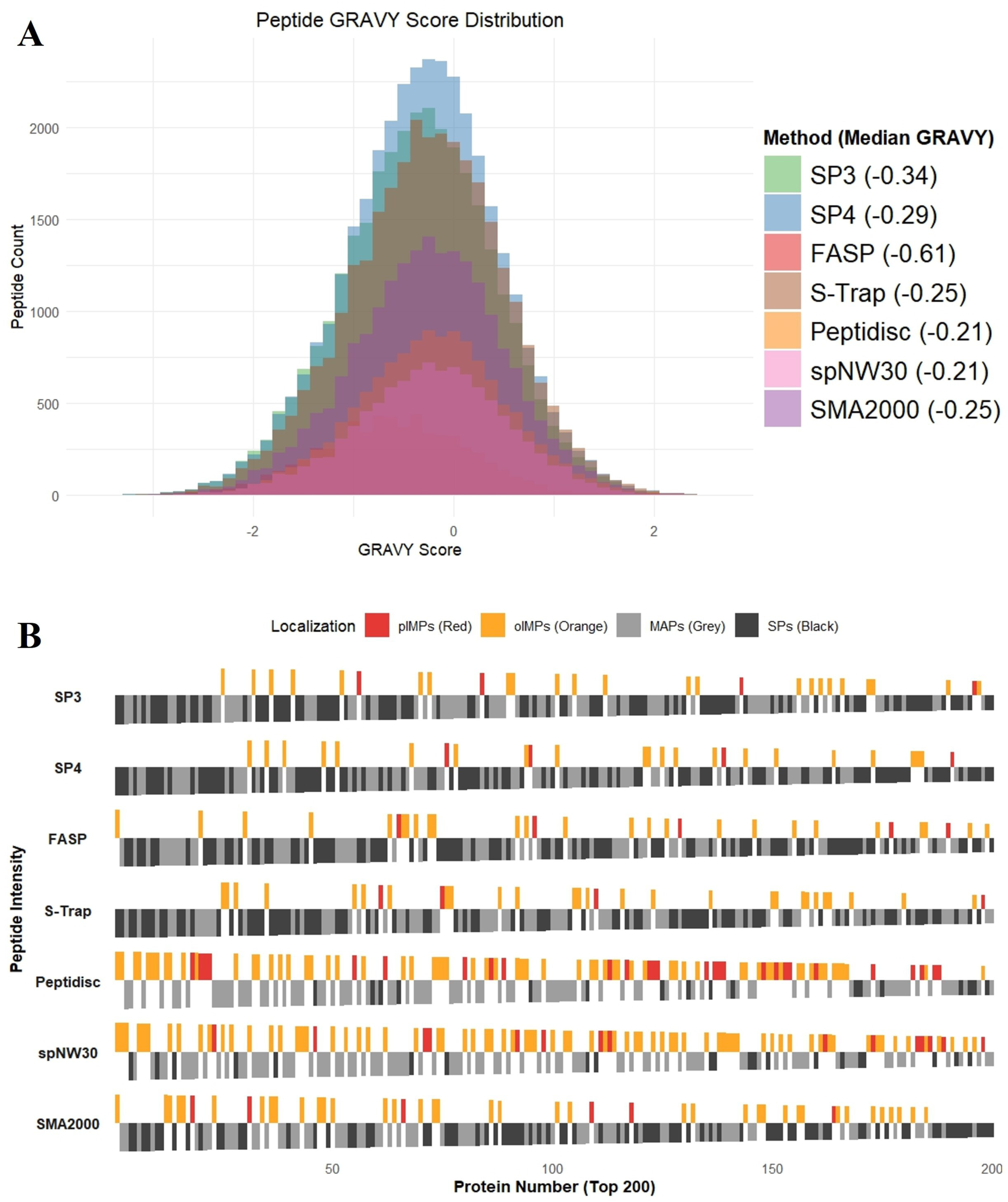
GRAVY Score across Methods and Enrichment Analysis of IMPs for the Top 200 Most Abundant Proteins **(A)** GRAVY scores were computed for each detected peptide across the workflows using the Kyte–Doolittle hydropathy scale, where positive values indicate hydrophobicity and negative values indicate hydrophilicity. Peptides were filtered to retain those with valid intensity values, and GRAVY scores were averaged per peptide sequence. Median GRAVY scores for each method are annotated in the legend. The distribution of GRAVY scores is shown as overlapping histograms, with individual methods colored distinctly. Peptides were derived from single-intensity replicate experiments and plotted using ggplot2 in R. **(B)** Bar plots showing the localization classification of the top 200 most enriched proteins (ranked by MaxLFQ intensity) from each workflow. Proteins are categorized into four groups based on annotation: pIMPs (red), oIMPs (orange), MAPs (grey), and SPs (black). The bar height. is scaled relative to rank to visually distinguish localization distribution along enrichment rank. Plotted using the ggplot2 R package.

**Figure 4.**
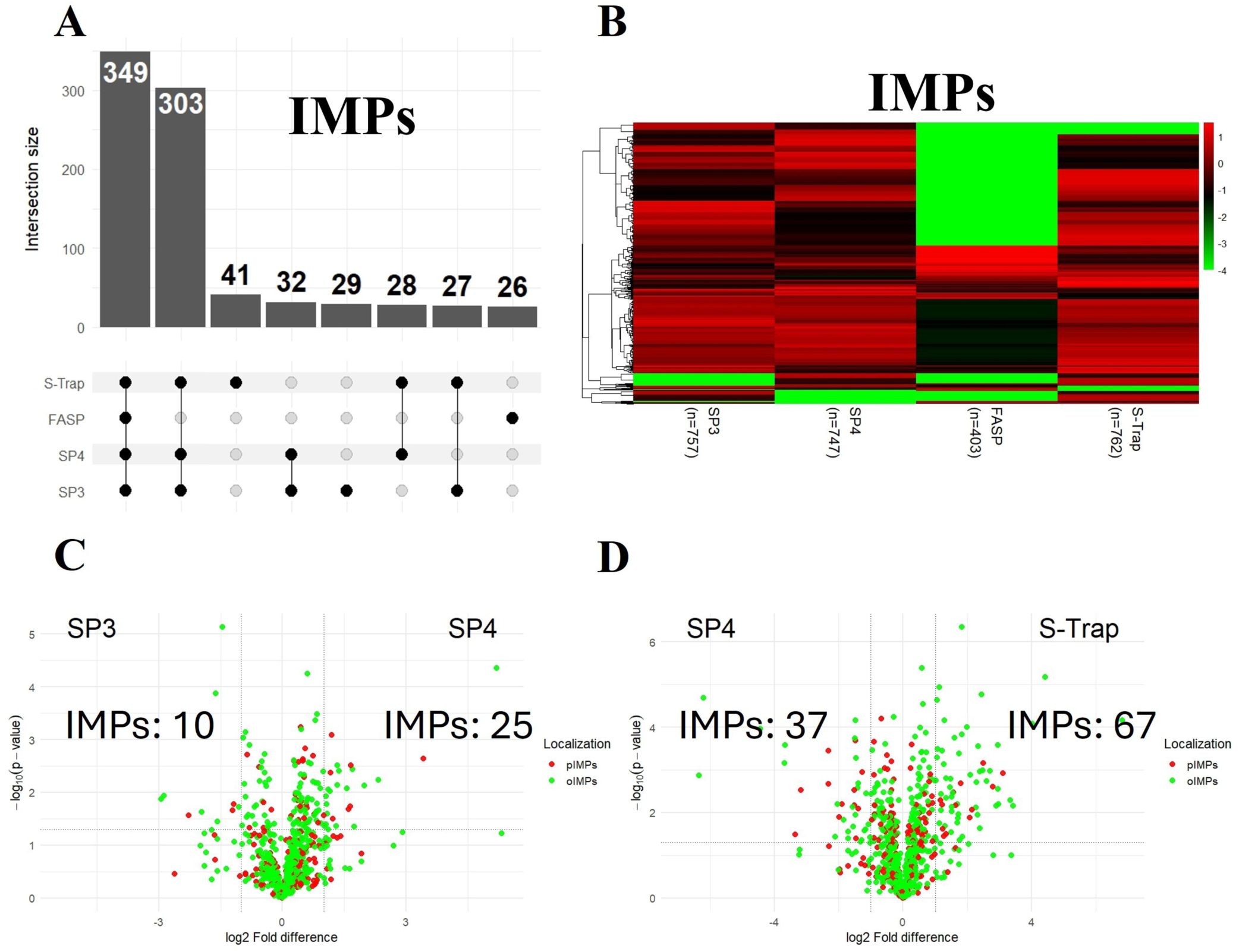
Comparative Recovery and Differential Enrichment Analysis of IMPs with the Solid-State Methods (A) UpSet plot showing the overlap of IMPs identified by SP3, SP4, FASP, and S-Trap. Intersections represent proteins shared between methods, with intersection sizes annotated above each bar. The analysis was performed in R using the Complex Upset package. **(B)** Heatmap plot showing the MaxLFQ intensity values of IMPs from the replicate with the highest number of protein identifications. Intensities were log₂-transformed and scaled row-wise. Missing values were imputed with a constant value of –4. The color scale represents standardized abundance values ranging from low (green) to high (red). Column labels indicate the number of IMPs detected per method. Heatmaps were generated using the pheatmap R package. (C-D) Volcano plots to illustrate protein enrichment differences between (C) SP3 and SP4, (D) SP4 and S-Trap. A two-tailed *t*-test was performed in Perseus with *s₀* = 0.1, and significance thresholds were set at log₂ FC ≥ ±1 and –log₁₀(p-value) ≥ 1.3. Transmembrane domains were predicted using Phobius to classify proteins as pIMPs (red) or oIMPs (orange). Only proteins with valid MaxLFQ intensity values in at least three out of six replicates were included, with missing values imputed using a width of 0.3 and a downshift of 1.8. Tables are included on either side indicating the number of differentially expressed IMPs that passed the p-value and log2 FC cutoff. Plots generated using the ggplot2 package in R.

**Figure 5.**
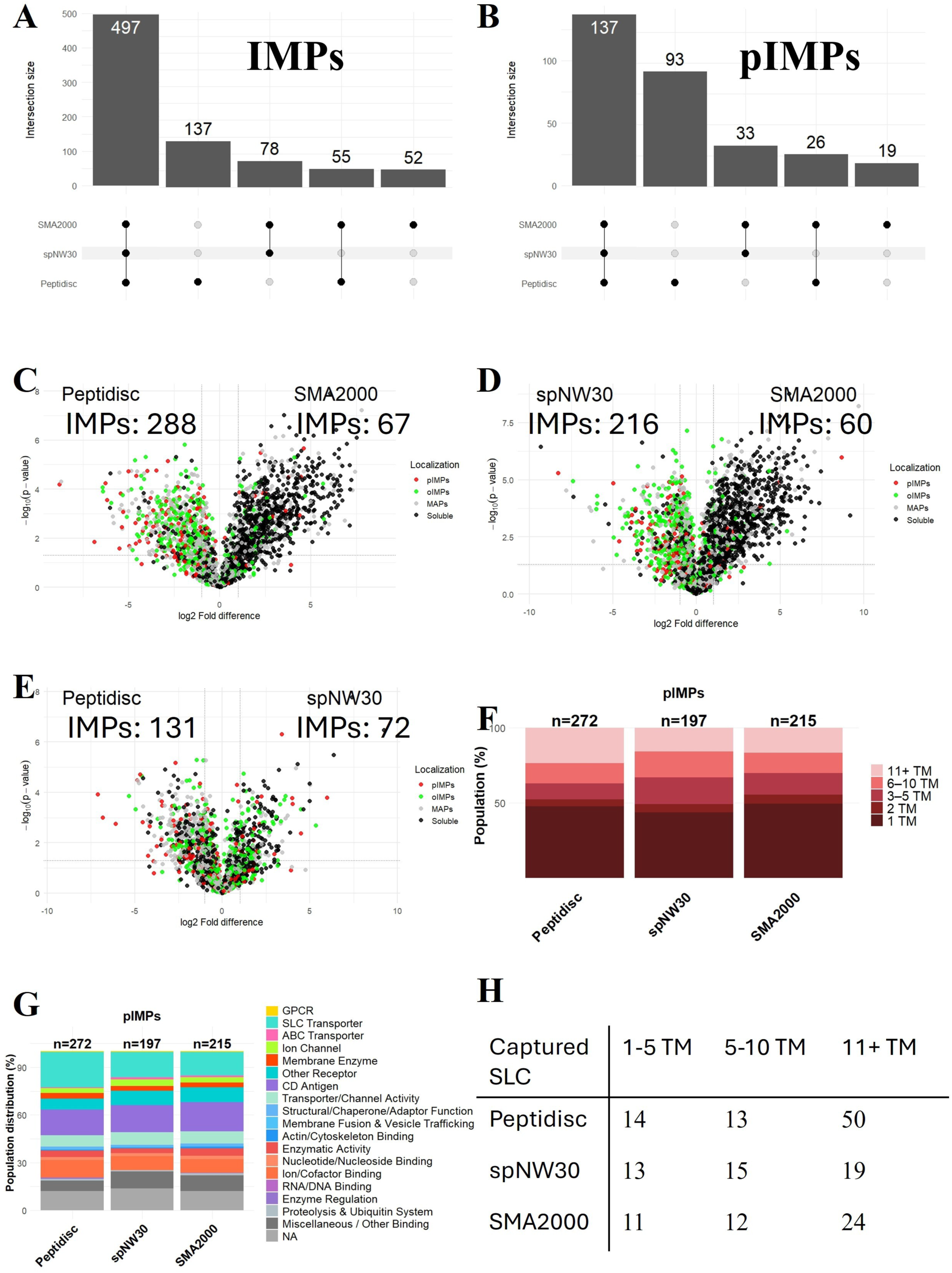
Comparative Recovery and Differential Enrichment Analysis of IMPs with the Membrane Mimetic Methods (A-B) UpSet plot showing the overlap of (A) IMPs and (B) pIMPs identified with Peptidisc, spNW30, and SMA2000. Intersections represent proteins shared between methods, with intersection sizes annotated above each bar. The analysis was performed in R using the ComplexUpset package. (C-E) Volcano plots illustrate protein enrichment differences between (C) Peptidisc and spNW30, (D) Peptidisc and SMA2000, (E) spNW30 and SMA2000. A two-tailed *t*-test was performed in Perseus with *s₀* = 0.1, and significance thresholds were set at log₂ FC ≥ ±1 and – log_₁₀_(p-value) ≥ 1.3. Transmembrane domains were predicted using Phobius to classify proteins as pIMPs: red, oIMPs: orange, MAPs and soluble proteins: gray. Only proteins with valid MaxLFQ intensity values in at least three out of six replicates were included, with missing values imputed using a width of 0.3 and a downshift of 1.8. Tables are included on either side indicating the number of differentially expressed IMPs, MAPs, or soluble proteins that passed the p-value and log2 FC cutoff. Plots generated using the ggplot2 package in R. **(F)** Stacked bar plot showing the relative distribution of transmembrane domain (TM) counts among pIMPs. TM counts were predicted by submitting protein FASTA sequences to Phobius. Proteins were grouped into five bins based on the number of predicted TM domains: 1 TM, 2 TM, 3–5 TM, 6–10 TM, and 11+ TM. The number of IMPs identified per method is indicated above each bar. Visualization was performed in R using the ggplot2 package. (G) Molecular pathway distribution of pIMPs categorized using keyword-matched terms from GO molecular function annotations. The number of proteins assigned per method is shown above each bar. Plots were generated using the ggplot2 package in R. (H) Table showing the count of SLC genes by the number of transmembrane segments.

**Figure 6.**
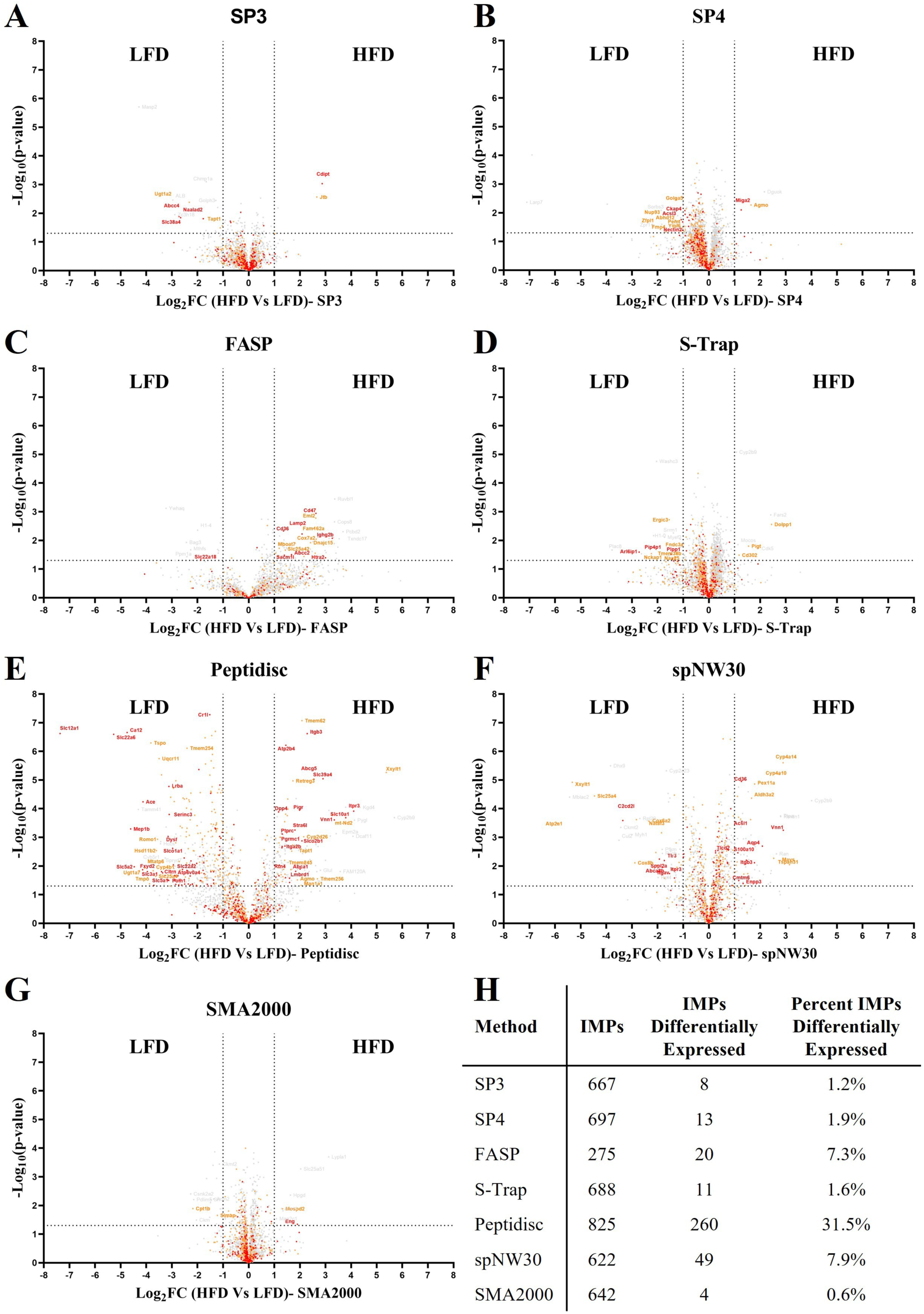
Capture of MP-Level Dysregulation between the LFD and HFD mLiver (A-G) Volcano plots of gene enrichment differences between LFD and HFD mLiver across (A) SP3, (B) SP4, (C) FASP (D) S-Trap, (E) Peptidisc, (F) spNW30, (G) SMA2000, with three biological replicates for the LFD and HFD. A two-tailed *t*-test was performed in Perseus with *s₀* = 0.1, and significance thresholds were set at log₂ FC ≥ ±1 and –log₁₀(p-value) ≥ 1.3. Transmembrane domains were predicted using Phobius to classify proteins as pIMPs: red, oIMPs: orange, MAPs and soluble proteins: gray. Only proteins with valid MaxLFQ intensity values in at least three out of six replicates were included, with missing values imputed using a width of 0.3 and a downshift of 1.8. Volcano plots were generated using GraphPad Prism 10. (H) Table summarizing the number and proportion of IMPs identified as differentially expressed.

**Figure 7.**
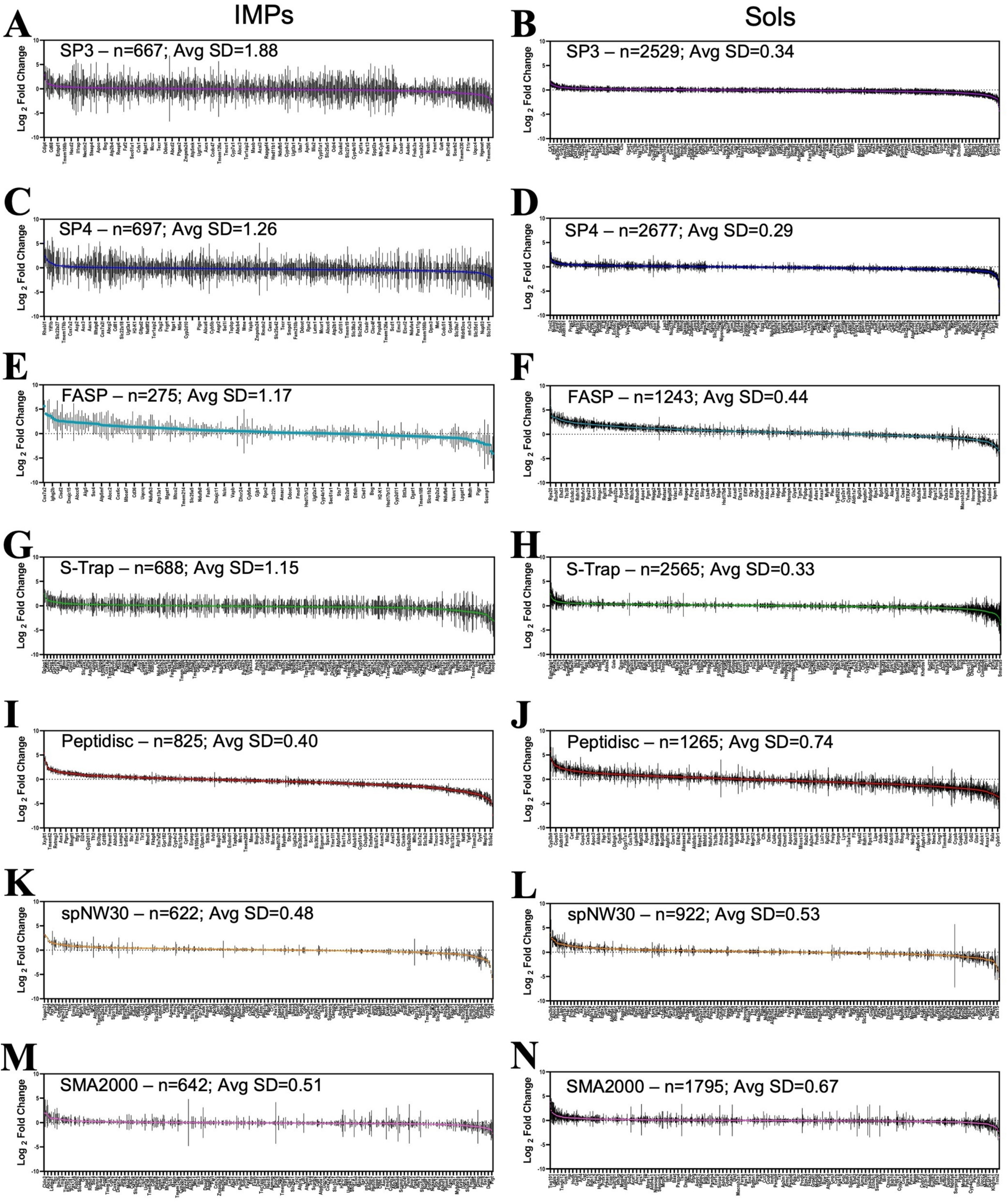
Standard Deviation in the Log2 FC for Protein Differentially Expressed Between the LFD and HFD mLiver Across Workflows (A-N) Plots display the standard deviation of log₂ FC in the Max LFQ Intensity between the LFD and HFD mLiver samples for (A-B) SP3, (C-D) SP4, (E-F) FASP, (G-H) S-Trap, (I-J) Peptidisc, (K-L) spNW30, (M-N) SMA2000. IMPs indicate either oIMPs or pIMPs, while Sols refer to MAPs + SPs. (The total n-number of proteins plotted and the average standard deviation of variance of log2 FC between all the proteins is shown). The total number of proteins plotted (n) and the average standard deviation (SD) of log2 FC obtained for each protein across three biological replicates for the LFD and HFD mLiver is shown. All plots were generated using GraphPad Prism 10.

**Figure 8.**
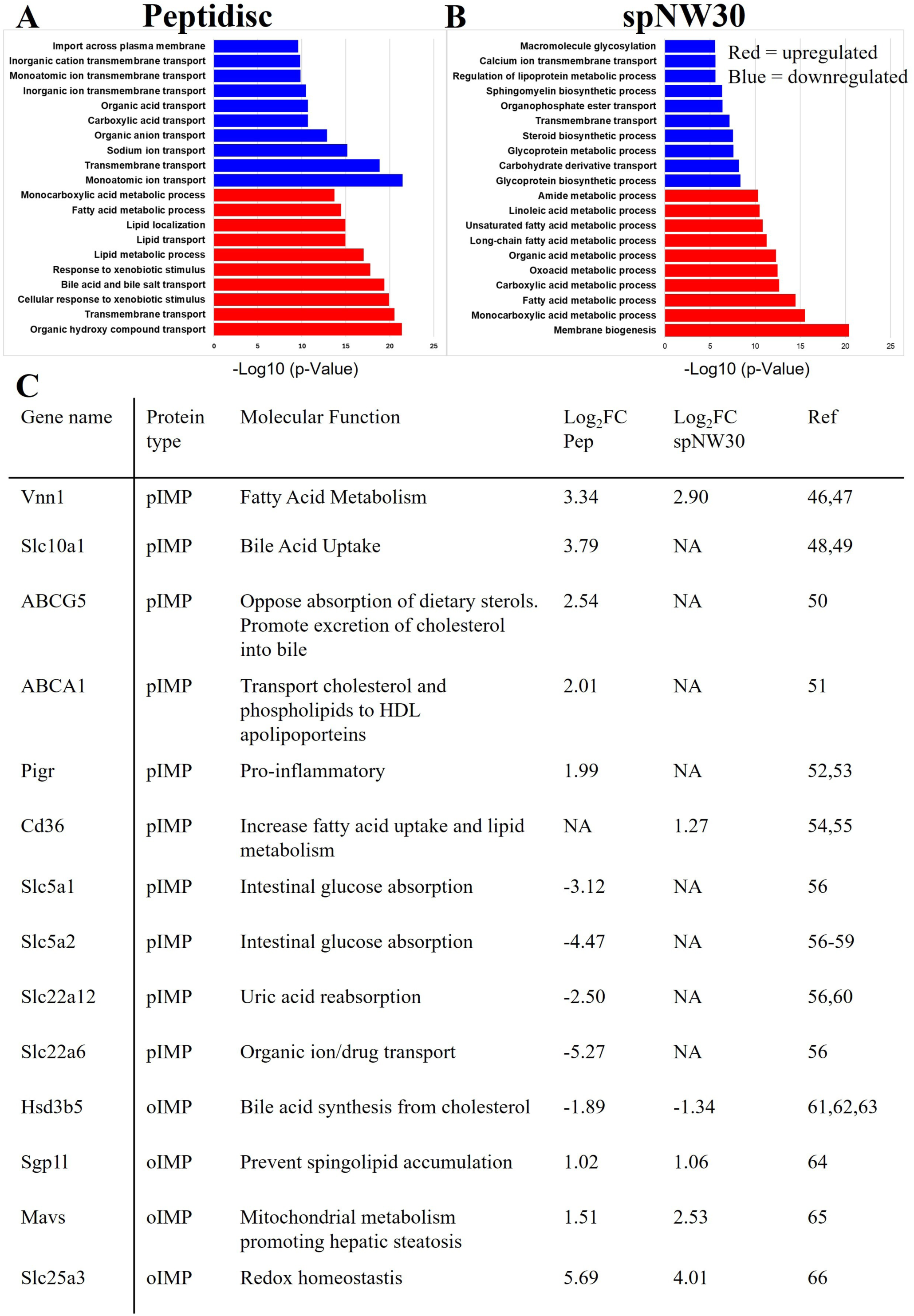
GO-Term Enrichment and Annotation for Differentially Expressed IMPs across Peptidisc and spNW30 in LFD and HFD mLiver. (A-B) The top 10 GO-term biological processes enriched among differentially expressed proteins in (A) Peptidisc and (B) spNW30 workflows, categorized as upregulated (red) or downregulated (blue) in the LFD vs. HFD mLiver, are presented. GO enrichment analysis was performed using DAVID Bioinformatics, and results were plotted with Excel. (C) Table summarizing the differential enrichment of genes associated with MASLD progression, as identified by Peptidisc and spNW30 in the LFD versus HFD comparison. Ref: Source publication as referenced in the Result section

#### Experimental Design and Statistical Rationale

The experimental design of the seven membrane proteomic workflows is presented in Figure 1. The overall strategy includes MP extraction, trypsin digestion, and quantitative LC-MS/MS analysis to benchmark each method’s performance. mLiver tissue samples were harvested from LFD and HFD cohorts. For each workflow—SP3, SP4, FASP, S-Trap, Peptidisc, SMA2000, and spNW30—three biological replicates per condition (n = 3 LFD, n = 3 HFD) were independently processed and analyzed to assess membrane proteome coverage, reproducibility, and the ability to detect differential protein expression.

Sample processing protocols varied by workflow, but all samples were ultimately subjected to trypsin digestion and desalting. Peptides were analyzed using LC-MS/MS in a data-dependent acquisition (DDA) mode (Figure 1H). Protein identification and quantification were performed using FragPipe v19.1 with MSFragger (v 4.1) and IonQuant (10 PPM precursor mass tolerance, 0.4 min retention time tolerance, 0.05 1/K0 ion mobility tolerance, top 3 ions). A FDR of 1% was applied at both the PSM and protein levels using target-decoy–based sequential filtering. The resulting combined_protein.tsv output was exported to Perseus (v1.6.15.0) for downstream statistical analysis.

Protein intensity and MaxLFQ values were log₂-transformed. Reproducibility between biological replicates was assessed using LFQ intensities and Pearson correlation coefficients. To determine differential protein abundance between LFD and HFD conditions, Student’s t-tests were performed with a permutation-based FDR correction (FDR < 0.05) and an artificial within-groups variance (s₀ = 0.1). Proteins were retained for analysis only if valid MaxLFQ intensity values were present in all replicates within both treatment groups. For missing values, intensity data were imputed from a normal distribution with a downshift of 1.8 standard deviations (SD) and a width of 0.3× SD relative to the sample distribution.

Proteins were considered significantly differentially expressed if they met a fold-change (FC) threshold of |log₂FC| ≥ 1 (corresponding to FC ≥ 2 or ≤ 0.5) and a p-value < 0.05 (–log₁₀(p) > 1.3). To assess physicochemical biases across workflows, GRAVY (Grand Average of Hydropathy) scores were calculated for all identified proteins using the Kyte–Doolittle scale. Pairwise comparisons of GRAVY score distributions between workflows were performed using the Wilcoxon rank-sum test with Bonferroni correction for multiple testing.

This experimental design enabled a rigorous and reproducible side-by-side comparison of proteomic workflows, providing insight into their relative strengths in enriching MPs and detecting protein-level dysregulation in a MASLD-relevant mLiver model.

## Results

### Workflow for Sample Preparation from mLiver Crude Membrane Extracts

To evaluate the performance of the seven proteomic workflows, we first isolated the crude membrane fraction from C57BL/6J mLiver. This fraction was prepared by homogenizing mLiver tissue with a French press in hypotonic buffer supplemented with a protease inhibitor and DNase. After low-speed centrifugation to remove nuclear and mitochondrial fraction, the supernatant was subjected to ultra-centrifugation (110,000 × *g*) to isolate the crude membrane fraction, which was subsequently used in all membrane proteomic workflows. Each workflow was performed in three replicates using liver tissue from three different male mice.

Prior to MS analysis, we conducted SDS-PAGE to assess the membrane proteomes obtained from each workflow (Fig. 2A). For the solid-phase proteomic workflows, an SDS-solubilized membrane extract served as the starting material; all workflows incorporated detergent removal either via bead-based adsorption or column-based trapping. This SDS extract exhibited broad protein banding from approximately 15 to over 250 kDa. The Peptidisc protein library was instead reconstituted from a DDM-solubilized membrane extract by reducing the detergent concentration below its critical micellar concentration and replacing it with the Peptidisc peptide during ultrafiltration. Following Ni-NTA affinity purification, this preparation displayed a wide range of protein bands spanning ∼5 to >250 kDa, along with an intense band at the bottom of the gel corresponding to the peptide scaffold. Similarly, the spNW30 library, also reconstituted from a DDM extract with the same protocol as Peptidisc, showed prominent bands above the 50 kDa marker, weaker bands below 50 kDa, and a dominant high-molecular-weight band corresponding to the MSP. Finally, the SMA library—prepared by direct solubilization of the native membrane using the SMA2000 copolymer—exhibited protein bands above 37 kDa, a broad smear below this marker, and a diffuse signal at the bottom of the gel attributable to the polymer.

In all cases, we trypsin-digested 100 µg of protein obtained from each workflow. For the solid-phase and SMA2000 workflows, the 100 µg input consisted of proteins entirely derived from the mLiver membrane fraction. In the spNW30 and Peptidisc workflows, the protein input included the scaffold components at a ratio relative to MP approximating to 1:1 and 1:2, respectively. Following digestion, equal amounts of the resulting peptide digests were injected for LC-MS/MS analysis.

### Replicate Consistency and Cross-Method Correlation

Before delving into detailed analysis of the MS data, we conducted a global assessment by examining intra-method reproducibility using Pearson correlation values across biological replicates. FASP showed high variability (r = 0.55–0.76), whereas the other six methods demonstrated strong reproducibility, with correlation values exceeding 0.8 (**Fig. S2**). To evaluate further the similarity between methods, we performed a pairwise correlation analysis using log₂-transformed Max LFQ intensities of shared proteins between workflows (**Fig. S3**). The resulting correlation r-values are presented Fig. 2B. SP3, SP4, and S-Trap exhibited strong inter-method correlation (r ≥ 0.95), while FASP showed weak correlation with all other methods (r = 0.22– 0.64). Peptidisc and spNW30 showed moderate correlation (r = 0.75) and weak correlation with all other methods (r = 0.22 to 0.68). SMA2000 showed intermediate correlation with both solid-phase (r = 0.87 for the SMA2000/SP3 pair) and membrane mimetic workflows (r = 0.68 for the SMA2000/spNW30 pair). These correlation patterns were further supported by principal component analysis (PCA), where SP3, SP4, and S-Trap clustered together, whereas FASP resolved separately. Peptidisc and spNW30 formed a distinct cluster and SMA2000 occupied an intermediate position between solid-phase and membrane mimetics (Fig. 2C).

### Hydrophobicity, Subcellular Localization, and Enrichment Patterns of IMPs across Methods

We first assessed the physicochemical properties of peptides obtained by each method, by comparing their GRAVY scores (Fig. 3A). Peptidisc and spNW30 exhibited the highest enrichment of hydrophobic peptides, reflected in a positive median score (both −0.21). SMA2000 and S-Trap showed intermediate hydrophobicity (both −0.25), whereas SP3, SP4 and FASP displayed higher overall hydrophilicity (−0.29, −034, and −061, respectively). Bonferroni-adjusted pairwise Wilcoxon tests revealed that Peptidisc and spNW30 did not differ significantly from each other, nor did S-Trap and SMA2000 (**Fig. S4**).

We next classified the list of total proteins (TPs) identified into five categories based on their GO-term annotation (**Table 1**): soluble proteins (SPs), membrane-associated proteins (MAPs), and integral membrane proteins (IMPs) when carrying at least one one predicted α-helical transmembrane segment. The IMPs were further subdivided based on predicted subcellular localization into plasma membrane (pIMPs) and other membrane compartments, including organelles (oIMPs). Among the solid-phase methods, SP4 identified the most TPs (3353 ± 42) while S-Trap identified the most IMPs (557 ±12). Among all methods, Peptidisc captured the highest number of IMPs (646 ± 17, accounting for 40% ± 0.4 of TPs).

**Table 1:** Number and Types of Proteins Captured Across the Workflows. The table summarizes the mean ± standard deviation of protein identifications across three biological replicates for each method. Reported categories include total proteins (TPs), soluble proteins (SPs), membrane-associated proteins (MAPs), and integral membrane proteins (IMPs), with IMPs further divided into plasma membrane (pIMPs) and organelle membrane (oIMPs) subsets. The table also includes the proportion (Ratio) of IMPs and pIMPs relative to the total proteome. Protein quantification was performed using LFQ values generated by MSFragger (v4.1) within FragPipe (v22.0) from the raw MS data. Transmembrane domains were predicted using Phobius, while protein localization (TP, SP, MAP, IMP, oIMP, or pIMP) was determined based on GO terms from the Cellular Component and Subcellular Localization categories. IMPs were defined as proteins with at least one predicted transmembrane segment and GO terms containing “Plasma Membrane” or organelle-specific membrane annotations (e.g., ER, Golgi, endosome, lysosome, exosome, peroxisome, mitochondria, or nuclear membrane). oIMPs are a subset of IMPs associated with organelle membranes, while MAPs were defined as proteins lacking transmembrane domains but annotated with the “Plasma Membrane” GO term.

|  | TPs |  |  | IMPs |  |  |  |  |
| --- | --- | --- | --- | --- | --- | --- | --- | --- |
| MS Sample | Total | Soluble | Membrane | Integral | Integral | Integral | Ratio | Ratio |
| (mean ± SD) | Proteins | Proteins | Associated | Membrane | Membrane | Membrane | IMPs / TPs | pIMPs / TPs |
|  |  |  | Proteins | Proteins | Proteins | Proteins |  |  |
|  |  |  |  |  | Plasma | Organelle |  |  |
|  | (TPs) | (SPs) | (MAPs) | (IMPs) | (pIMPs) | (oIMPs) | % IMPs | % pIMPs |
| SP3 | 3317±74 | 1629±34 | 1144±26 | 543±14 | 241±5 | 502±10 | 16±0.10 | 7±0.04 |
| SP4 | 3353±42 | 1664±20 | 1153±14 | 536±10 | 240±3 | 496±9 | 16±0.13 | 7±0.06 |
| FASP | 1996±364 | 953±166 | 663±128 | 379±70 | 113±27 | 266±43 | 19±0.14 | 6±0.31 |
| S-Trap | 3332±12 | 1616±6 | 1158±8 | 557±12 | 241±5 | 517±8 | 17±0.33 | 7±0.12 |
| Peptidisc | 1886±35 | 483±13 | 649±31 | 753±42 | 276±32 | 477±13 | 40±0.55 | 15±0.50 |
| spNW30 | 1598±29 | 451±16 | 506±7 | 642±10 | 194±4 | 448±7 | 40±0.46 | 12±0.16 |
| SMA2000 | 2511±28 | 1017±11 | 819±7 | 675±11 | 212±4 | 463±6 | 27±0.13 | 8±0.09 |

We annotated using this classification the top 200 most abundant proteins identified in each workflow. Peptidisc (92) and spNW30 (88) exhibited the highest enrichment of IMPs, followed by SMA2000 (44), while the solid-phase methods showed much lower enrichment levels (25–31) (Fig. 3B). This pattern was further supported by quartile-based analysis of the full dataset. In the top abundance quartile, Peptidisc again showed the highest IMP representation (47.7%), closely followed by spNW30 (46.3%), then SMA2000 (25.4%), whereas the solid-phase methods remained comparatively low (14.7–17.9%) (**Fig. S5**).

Together, these findings highlight distinct hydrophobicity and IMP enrichment profiles across the seven workflows. Peptidisc and spNW30 workflows demonstrated the strongest enrichment of hydrophobic peptides and IMPs. SMA2000 showed intermediate hydrophobicity and IMP enrichment, while solid-phase methods (SP3, SP4, S-Trap) and FASP identified more hydrophilic peptides and exhibited lower IMP representation.

### IMPs Recovery with the Solid-State Methods

We performed an UpSet plot analysis to assess the overlap and uniqueness of IMP proteins identified by each of the solid-phase extraction methods. The largest group comprised 349 IMPs consistently captured by all four methods, followed by 303 IMPs shared among SP3, SP4, and S-Trap, and 41 IMPs uniquely identified by S-Trap (Fig. 4A). These results suggest that a substantial subset of IMPs is commonly enriched by SP3, SP4, and S-Trap but not effectively captured by FASP. To explore this further, we conducted a heatmap analysis using log₂-transformed MaxLFQ intensity values for IMPs identified across the four methods. This side-by-side comparison provided additional insight into the degree of overlap and relative protein enrichment across datasets. Consistent with the UpSet plot findings, FASP did not yield a distinct or unique proteome profile, but instead recovered a smaller subset of proteins also detected by the other three solid-phase methods (Fig. 4B).

A volcano plot analysis was used to assess the relative enrichment of proteins shared between workflows. For each protein detected in both workflows, the log₂-transformed MaxLFQ intensity values from three biological replicates per workflow (six replicates total) were compared. Proteins were considered differentially enriched if they exhibited a log₂ FC ≥ 1 or ≤ – 1 and a –log₁₀ p-value > 1.3, where the p-value reflects the consistency of intensity differences across replicates, higher variability yields lower statistical significance. This analysis highlights proteins with large and reproducible differences in abundance between workflows. SP4 enriched more IMPs compared to SP3 (25 vs. 10; Fig. 4C), while S-Trap enriched a higher number of IMPs compared to SP4 (67 vs. 37; Fig. 4D). Together, SP3, SP4, and S-Trap captured overlapping protein libraries, however S-Trap exhibited the highest relative recovery of IMPs among the three methods.

### IMPs Recovery with Membrane Mimetics

We applied an UpSet plot analysis to assess the overlap and uniqueness of IMPs captured by the membrane mimetic workflows. A total of 497 IMPs were consistently identified across all three methods, while 137 IMPs were uniquely captured by the Peptidisc workflow (Fig. 5A). A similar trend was observed for pIMPs, with 137 shared among all methods and 93 uniquely identified by Peptidisc (Fig. 5B). Notably, the uniquely captured pIMPs represented 68% of the total shared pIMPs, whereas uniquely captured IMPs accounted for only 28% of the shared IMPs. These findings suggest that the Peptidisc workflow provides preferential enrichment for pIMPs compared to the other two membrane mimetic methods.

A volcano plot analysis was also conducted to compare the enrichment of shared proteins across the membrane mimetic workflows (Fig. 5C, **5D** and **5E**). Peptidisc demonstrated greater enrichment of IMPs compared to SMA2000, capturing 288 versus 67 proteins, respectively.

Similarly, spNW30 also outperformed SMA2000, with 216 IMPs identified versus 60. In contrast, SMA2000 showed stronger enrichment of soluble proteins relative to both Peptidisc and spNW30 (288 vs. 67 in Peptidisc; 216 vs. 60 in spNW30). When directly comparing Peptidisc and spNW30, Peptidisc enriched a higher number of IMPs (131 vs. 72 in spNW30). These results indicate that SMA2000 preferentially enriches soluble proteins, whereas both Peptidisc and spNW30 show enhanced capture of IMPs, with Peptidisc demonstrating the strongest enrichment overall.

### Characterization of pIMPs Captured by Membrane Mimetic Workflows

The observation that Peptidisc captures a substantial subset of pIMPs that are either underrepresented or undetected by spNW30 or SMA2000 prompted further evaluation. For instance, mass distribution analysis reported that Peptidisc enriched for pIMPs with larger molecular weight (>70 kDa) compared to SMA2000 and spNW30 (43% vs. 38%; **Fig. S6**). Furthermore, Peptidisc demonstrated superior enrichment for proteins with 11 or more transmembrane segments, capturing 80 compared to 31 by spNW30 and 37 by SMA2000 (Fig. 5F). Functional categorization based on GO molecular function terms showed that Peptidisc also captured a greater number of solute carrier (SLC) transporters, identifying 81 compared to 31 for spNW30 and 34 for SMA2000 (Fig. 5G). Consistent with this, UpSet plot analysis revealed that 45 SLC transporters were shared across all three workflows, but Peptidisc uniquely captured 37, compared to just 5 and 3 uniquely identified by SMA2000 and spNW30, respectively **(Fig. S7A**). Furthermore SLC transporters with 11 or more transmembrane segments comprised 76% of uniquely identified SLCs for Peptidiscs (**Fig. S7B**) Notably, all three membrane mimetic workflows captured similar numbers of SLC transporters with 1–10 transmembrane segments, however Peptidisc detected considerably more with 11 or more segments (Peptidisc: 50; SMA2000: 24; spNW30: 19; Fig. 5H).

Together, these findings suggest that Peptidisc’s preferential capture of pIMPs is driven by its enrichment of proteins with higher transmembrane segment counts and consequently larger molecular weights. This trend is consistent with the increased number of SLC transporters identified by Peptidisc, particularly those with 11 or more transmembrane segments, compared to SMA2000 and spNW30. Notably, many SLC transporters form higher-order oligomeric structures, such as heterodimers and heterotetramers(45), which increase their overall hydrophobicity and structural complexity in the native membrane. These features may enhance compatibility with the flexible, amphipathic Peptidisc scaffold, compared to the more rigid architecture of spNW30.

### Application of the Workflows to LFD and HFD mLivers

To evaluate the ability of the seven workflows to detect biologically relevant changes in the membrane proteome, we performed a comparative analysis using healthy and diseased mLiver tissue from C57BL/6J mice fed either a LFD (10% kcal from fat) or a HFD (60% kcal from fat). Following MS analysis, a moderate decrease in TP count was observed under HFD conditions, with an average of 98.7 proteins detected (**Table 2**). We performed a differential protein enrichment analysis with volcano plots to compare MP expression between LFD and HFD mLiver samples (Fig. 6). Overall, SP3, SP4, FASP, S-Trap, and SMA2000 detected relatively few differentially expressed IMPs (ranging from 4 to 20), whereas spNW30 identified a moderate number (n = 49). In sharp contrast, Peptidisc detected a substantially larger set (n = 260).

**Table 2.** Number and Types of Proteins Captured in the LFD and HFD mLiver Across the Workflows. The table reports the mean ± standard deviation of protein identifications across three biological replicates for each proteomic workflow applied to either LFD or HFD mLiver tissue. Quantified categories include total proteins (TPs), soluble proteins (SPs), membrane-associated proteins (MAPs), and integral membrane proteins (IMPs). IMPs are further divided into plasma membrane (pIMPs) and organelle membrane (oIMPs) subsets. The percentage of IMPs and pIMPs relative to the total proteome is also shown. Label-free quantification (LFQ) values were generated using MSFragger (v4.1) within FragPipe (v22.0) from raw MS data. Transmembrane domains were predicted using Phobius. Protein subcellular localization (TP, SP, MAP, IMP, oIMP, pIMP) was determined using GO terms from the Cellular Component and Subcellular Localization categories. IMPs were defined as proteins containing at least one predicted transmembrane segment and annotated with GO terms indicating plasma or organelle membranes. oIMPs represent the subset of IMPs localized to organelle membranes (e.g., ER, Golgi, mitochondria, vesicles, and nucleus). MAPs were defined as proteins lacking transmembrane domains but annotated with the “Plasma Membrane” GO term.

| MS Sample<br>(mean ± SD) | TPs | | | IMPs | | | | |
| --- | --- | --- | --- | --- | --- | --- | --- | --- |
|  | Total | Soluble | Membrane | Integral | Integral | Integral | Ratio | Ratio |
|  | Proteins | Proteins | Associated | Membrane | Membrane | Membrane | IMPs / TPs | pIMPs / |
|  |  |  | Proteins | Proteins | Proteins | Proteins |  | TPs |
|  |  |  |  |  | Plasma | Organelle |  |  |
|  | (TPs) | (SPs) | (MAPs) | (IMPs) | (pIMPs) | (oIMPs) | % IMPs | %pIMPs |
| SP3 LFD | 3317±74 | 1629±34 | 1144±26 | 543±14 | 241±5 | 502±10 | 16±0.10 | 7±0.04 |
| SP3 HFD | 3221±147 | 1581±66 | 1123±49 | 517±33 | 232±9 | 485±24 | 16±0.30 | 7±0.09 |
| SP4 LFD | 3353±42 | 1664±20 | 1153±14 | 536±10 | 240±3 | 496±9 | 16±0.13 | 7±0.06 |
| SP4 HFD | 3303±38 | 1630±17 | 1144±14 | 529±9 | 239±4 | 491±8 | 16±0.12 | 7±0.13 |
| FASP LFD | 1996±364 | 953±166 | 663±128 | 379±70 | 113±27 | 266±43 | 19±0.14 | 6±0.31 |
| FASP HFD | 1876±117 | 894±46 | 617±42 | 365±30 | 107±8 | 258±22 | 19±0.37 | 6±0.14 |
| S-Trap LFD | 3332±12 | 1616±6 | 1158±8 | 557±12 | 241±5 | 517±8 | 17±0.33 | 7±0.12 |
| S-Trap HFD | 3128±118 | 1524±44 | 1107±39 | 497±36 | 217±10 | 480±26 | 16±0.54 | 7±0.05 |
| Peptidisc LFD | 1886±35 | 483±13 | 649±31 | 753±42 | 276±32 | 477±13 | 40±0.55 | 15±0.50 |
| Peptidisc HFD | 1738±36 | 498±22 | 577±3 | 577±18 | 229±7 | 434±12 | 38±0.39 | 13±0.12 |
| spNW30 LFD | 1598±29 | 451±16 | 506±7 | 642±10 | 194±4 | 448±7 | 40±0.46 | 12±0.16 |
| spNW30 HFD | 1535±29 | 443±19 | 479±9 | 613±2 | 191±2 | 422±4 | 39±1 | 12±0.37 |
| SMA2000 LFD | 2511±28 | 1017±11 | 819±7 | 675±11 | 212±4 | 463±6 | 27±0.13 | 8±0.09 |
| SMA2000 HFD | 2501±28 | 1012±12 | 818±11 | 671±5 | 211±2 | 460±8 | 27±0.09 | 8±0.19 |

### Assessment of Method Accuracy in Capturing MP-Level Dysregulation

Before analysis of the dysregulated proteins, we first evaluated the accuracy of each workflow for capturing proteome dysregulation by assessing the variance in protein-level log₂ FC values across three biological replicates, separately for IMPs and soluble proteins (SPs + MAPs). This was motivated by our observation that membrane mimetic methods yielded robust detection of differentially expressed proteins in volcano plots, whereas solid-phase approaches such as SP3, SP4, FASP, and S-Trap detected very few.

We calculated the log₂ FC for each protein across three biological replicates and visualized the variance in these values using SD plots. To assess overall accuracy, we then computed the average SD across all proteins within each workflow. Solid-phase methods exhibited substantial variability for IMPs, with an average SD ranging from 1.15-1.88. In contrast, these workflows showed low variance for soluble proteins, with SP4 achieving the lowest average SD (0.29). On the other hand, membrane mimetic methods demonstrated markedly lower variability for IMPs, with average SDs ranging from 0.40 to 0.51. Notably, Peptidisc achieved the lowest SD (0.40) among all workflows, consistent with its enhanced sensitivity in detecting differentially expressed MPs (Fig. 7). This improved accuracy is likely attributable to the detergent-free nature of these workflows, the stabilization of MPs via amphipathic scaffolds, and the use of a Ni-NTA enrichment step. Collectively, these findings indicate that membrane mimetics, particularly Peptidisc, offer superior accuracy and are better suited than solid-phase methods for capturing MP-level dysregulation.

### Functional Annotation of Identified MP Dysregulation

We next conducted GO biological process enrichment analysis for differentially expressed IMPs captured by Peptidisc and spNW30 (Fig. 8A and Fig. 8B). Peptidisc-captured IMPs showed significant enrichment for MASLD-relevant pathways, including bile acid transport, lipid metabolism, and xenobiotic response, along with downregulation of various ion transport processes. In contrast, IMPs captured by spNW30 were enriched for processes related to membrane biogenesis and glycoprotein biosynthesis, with downregulation observed in sphingomyelin and calcium ion transport pathways. Notably, the average –log₁₀(p-value) of enriched pathways was higher for Peptidisc in both the upregulated (17.9 vs. 12.0 for spNW30) and downregulated sets (13.7 vs. 7.4 for spNW30), indicating greater statistical confidence in the enrichment results. These data suggest that Peptidisc identifies a broader range of dysregulated MPs associated with MASLD-related pathways compared to spNW30, leading to stronger and more statistically robust biological insights.

Focusing on IMPs previously known to be implicated in MASLD, several were captured as differentially expressed by both Peptidisc and spNW30 (*Vnn1*, *Hsd3b5*, *Sgp1l*, *Mavs*).

However, additional key IMPs were uniquely identified as differentially expressed by Peptidisc, including *Slc10A1, Slc5a1, Slc5a2, Slc22a12, Slc22a6, Slc25a3, ABCG5, ABCA1*, and *Pigr*, while only *Cd36* was uniquely detected by spNW30 (Fig. 8C). These proteins participate in diverse pathways reportedly related to MASLD (46–66), suggesting that Peptidisc captures a broader MP-level dysregulation than being limited to specific biological processes. Collectively, these findings strongly suggest that Peptidisc offers superior accuracy, sensitivity, and biological relevance for detecting MP-level dysregulation in the context of MASLD, outperforming spNW30 across multiple evaluation criteria.

## Discussion

Despite their critical roles in biology and disease, MPs remain difficult to analyze using standard proteomic approaches(15). In this study, we present the first comprehensive, head-to-head comparison of seven leading workflows designed to resolve the membrane proteome. Our evaluation revealed a fundamental trade-off between two dominant approaches: solid-phase extraction methods identify a greater number of TPs, while membrane mimetic strategies detect fewer TPs but achieve superior recovery and proportional enrichment of IMPs. This outcome aligns with the design of membrane mimetic workflows, which are specifically optimized to stabilize hydrophobic, transmembrane segments–containing proteins and incorporate Ni-NTA– based depletion of soluble contaminants. In the sections below, we highlight key mechanistic factors that could explain their differing performance profiles.

Among the solid-phase extraction methods, FASP demonstrated the lowest overall performance, yielding the fewest TPs and IMPs, along with high variability. This inefficiency, also observed by others(67,68), stems from its reliance on protein aggregation on a filter—a process particularly problematic for MPs due to their hydrophobic nature, which promotes irreversible clumping and subsequent protein loss. These limitations have prompted the development of modified FASP protocols using sodium deoxycholate instead of urea and preconditioning filters with 5% Tween-20 to mitigate aggregation(68), however, the effectiveness of these adjustments remains uncertain.

To limit irreversible protein aggregation, SP3 and SP4 protocols instead use surface-based capture. In our hands, SP4 showed greater enrichment of hydrophobic peptides compared to SP3, as indicated by GRAVY score analysis. This observation is consistent with a previously reported underlying mechanism(69): SP3 captures proteins through a combination of hydrophilic interactions with carboxyl-group coated beads, leading to preferential retention of hydrophilic proteins. SP4 instead primarily relies on protein aggregation onto silica glass beads. These beads are capable of hydrogen bonding with polar residues, but the resulting hydrophilic interactions are weaker than with the carboxylated beads used in SP3, contributing to SP4’s relative enrichment of hydrophobic proteins.

Among the solid-phase approaches, S-Trap emerged as the top performer, capturing the highest number of IMPs and hydrophobic peptides with the lowest variability. This enhanced performance is likely stemming from the acidification step unique to the S-Trap protocol, which protonates amino acid side chains, leading to protein unfolding and exposure of hydrophobic regions. Unlike SP3 and SP4, which depend on interactions with bead chemistry, S-Trap rather promotes a broader binding of hydrophobic regions onto filters. To our knowledge, this is the first study to show that S-Trap preferentially enriches hydrophobic proteins relative to SP3, SP4, and FASP, establishing it as a strong candidate for membrane proteome analysis. However, S-Trap use of proprietary spin filters renders it more expensive than other methods. In contrast, SP4 uses readily available reagents, making it the most cost-effective among the solid-phase options without sacrificing MP capture efficiency. Importantly, all solid-phase methods exhibited lower accuracy in detecting MP-level dysregulation between the LFD and HFD liver samples compared to membrane mimetic strategies. This reduced performance is likely due to the multiple sample processing steps, such as detergent removal, protein precipitation, centrifugation, and urea-based resuspension, that together introduce variability across replicates.

Shifting focus to membrane mimetic strategies, we evaluated SMA2000, a widely used polymer-based system. Our analysis builds on the workflow introduced by Brown et al, who compared various SMALP formulations for profiling the membrane proteome in human embryonic kidney 293 cells(34). Their reported identification of 1,725–1,931 proteins across different SMA variants is consistent with the 2,511 proteins identified in our study. Interestingly, despite being a bona fide membrane mimetic, SMA2000 produced a proteome profile more closely aligned with solid-phase methods. Specifically, it enriched peptides with hydrophobicity comparable to those captured by S-Trap but exhibited a lower accuracy in capturing differences between the LFD and HFD compared to other mimetic workflows. Following Brown et al., we used the MTBE precipitation step in the workflow to separate proteins from the excess SMA polymer and lipids. This step reduces LC-MS/MS column contamination and signal suppression from the negatively charged polymer. This step, however, may also promote protein aggregation and loss during processing. contributing to SMA higher variability. Additionally, the SMA2000 workflow detected a lower proportion of IMPs compared to Peptidisc and spNW30, suggesting that the higher abundance of soluble peptides in its dataset may overshadow IMP detection and contribute to its reduced accuracy in capturing MP-level dysregulation. Furthermore, unlike spNW30 and Peptidisc, the SMA protocol lacks a Ni-NTA–based enrichment step, which may further limit its performance. Overall, while SMA2000 demonstrates promising features and reliable protein identification, its utility could be significantly enhanced by future refinements, such as improving MS compatibility of the polymer or integrating affinity purification steps.

To further dissect membrane mimetic strategies, we compared Peptidisc and spNW30, two platforms that both employ amphipathic scaffolds in combination with Ni-NTA purification to enrich MPs. Despite their conceptual similarity and shared use of reconstitution protocols, Peptidisc consistently outperformed spNW30 in both detection and enrichment of IMPs, particularly those with multiple transmembrane helices. Furthermore, when applied to compare membrane proteomes from murine livers under LFD versus HFD conditions, Peptidisc detected the highest number of differentially expressed IMPs and exhibited the lowest standard deviation in log₂ FC, indicating that Peptidisc is more accurate in detecting MP-level dysregulation.

Accordingly, the IMPs differentially regulated by Peptidisc showed stronger statistical enrichment in pathways associated with MASLD compared to those captured by spNW30. We attribute these divergences to fundamental differences in the self-assembly mechanisms underlying each platform.

Peptidisc employs short, synthetic amphipathic peptides that spontaneously and flexibly self-assemble around solubilized MPs during detergent removal. This mode of assembly, defined as a “one-size-fits-all” (35), allows the scaffold to adapt to the shape and topology of the embedded protein, enabling efficient encapsulation of diverse MPs regardless of their topology. The simplicity of this approach is likely to result in a more reproducible and accurate outcome, since it applies minimal constraints on protein size, structural complexity, or oligomerization state.

In contrast, the spNW30 system relies on the incorporation of IMPs in a fixed ∼30 nm discoidal lipid-bilayer structure supported by two large MSPs(31). This assembly process, inherently stochastic, requires the precise incorporation of both phospholipids and target proteins, making it difficult to control, especially when generating complex libraries of diverse MPs from crude detergent extracts. While the disc size (∼30 nm) should, in principle, accommodate large MPs, coarse-grained molecular dynamics simulations have shown that nanodiscs form through disordered protein–lipid intermediates and may adopt non-ideal conformations, such as incomplete double-belt structures or single MSPs folding back on themselves(70), which may hinder the proper incorporation of larger or structurally complex proteins. These factors together may introduce batch variability and affect reproducibility.

Moreover, because the spNW30 workflow was applied to mLiver crude membranes, the captured IMPs were already embedded in endogenous lipids, and no exogenous lipids were added during reconstitution. As such, no lipid optimization step was performed, which may have limited the efficiency of IMP reconstitution in the spNW30 nanodiscs(71).

The reduced performance of spNW30 may also be explained by practical and biochemical limitations during MS analysis. MSPs are large recombinant proteins that generate a substantial number of tryptic peptides upon digestion, increasing background signal and potentially obscuring the detection of low-abundance MPs. Furthermore, because MSPs are expressed and purified from *E. coli*, host-cell proteins do persist in the final nanodisc preparation. As the MS data were processed using a mouse reference proteome, these *E. coli*-derived peptides could be misassigned as mouse proteins, leading to inaccurate identifications. In contrast, the Peptidisc scaffold is chemically synthesized and free from biological contaminants, minimizing background interference in proteomic analyses. Moreover, spiking the Peptidisc peptide sequence into the mouse reference database ensures their exclusion from the target protein dataset.

Together, these results establish Peptidisc as the most effective of the seven workflows evaluated in this study, with the highest IMP capture, sensitive and reproducible detection of differential expression. Its low technical variability and simple workflow render it well suited for membrane proteome profiling, particularly in comparative or differential expression studies.

Looking ahead, our findings highlight a promising opportunity to expand membrane proteome coverage through the strategic integration of complementary workflows. For example, combining membrane mimetics with distinct enrichment profiles—such as Peptidisc and Nanodisc—could enable broader and more balanced recovery of diverse MP classes. The modular design of nanodiscs, in particular, allows tunable enrichment using panels with varying diameters (e.g., spNW15, 50, 80, 100)(31). Incorporating tailored exogenous lipids during reconstitution with mLiver membranes may further enhance compatibility with mammalian proteins and improve capture efficiency(71). In parallel, advancing the SMALP copolymer to support affinity enrichment or direct compatibility with MS could offer an orthogonal profiling strategy, improving the detection of differentially expressed MPs.

Beyond protein identification and quantification, our comparative analysis calls for a broader implementation of membrane mimetic workflows. For instance, the Peptidisc was recently shown to enable efficient thermal proteome profiling for MS-based ligand detection(72). Similarly, the SMA system was applied in yeast display–based nanobody discovery(73), suggesting that comparative evaluation of membrane mimetics could inform nanobody screening strategies. Together, these new applications underscore the importance of integrating proteomic benchmarking with functional validation to guide workflow selection for structural, biochemical, and therapeutic applications.

This study presents the first comprehensive evaluation of four solid-phase (SP3, SP4, FASP, S-Trap) and three membrane mimetic (Peptidisc, spNW30, SMA2000) workflows for profiling the LFD and HFD mLiver membrane proteome. We found that the solid-phase methods yield higher total protein identifications, while membrane mimetics provide superior enrichment of IMPs. Among these, Peptidisc stands out for its capture and enrichment of pIMPs, particularly those with 11 or more transmembrane segments. Although the SMA2000 workflow involves the use of a membrane mimetic polymer, its proteomic behavior was intermediate, showing stronger correlation with solid-phase workflows. When assessing protein-level dysregulation between LFD and HFD conditions, Peptidisc clearly outperformed the other six workflows, identifying the highest number of differentially expressed IMPs which are proteins in pathways that would be expected to be implicated in MASLD and showing the highest accuracy in capturing the MP dysfunction. Together, these findings provide a practical framework for selecting membrane proteomic workflows based on experimental goals such as protein localization, total protein yield, MP enrichment, and the capture of MP-level dysfunction.

## Data availability

The mass spectrometry proteomics data have been deposited to the ProteomeXchange Consortium via the PRIDE (74) partner repository with the dataset identifier PXD067014. Information related to the mass spectrometry data can be found in the Supplemental Data. Reviewer access details: Log in to the PRIDE website using the following details: Project accession: PXD067014, Token: 1BrcYXFAN18k. Alternatively, reviewer can access the dataset by logging in to the PRIDE website using the following account details: Username: Password: kiucE1Wxso4N

## Supplemental data

This article contains supplemental data.

## Acknowledgements

Work in the Duong lab was supported by the CIHR Project grant PG20R34019. Work in the Babu lab was supported by the Canada Foundation for Innovation and CIHR Foundation grant FDN-154318. Work in the Al Batran lab was supported by CIHR Project grant PJT-195730. R.A.B. is a Research Scholar with the Fonds de Recherche du Québec - Santé (FRQS) and a New Investigator for the Heart and Stroke Foundation of Canada (HSF). A.B. is supported by a UBC four-year fellowship.

## Author Contributions

Conceptualization: F.A., F.D.v.H. Methodology: F.A.

Validation: F.A.

Formal Analysis: F.A., A.B.

Investigation: F.A., H.A., R.S.J., A.M.A.

Resources: F.A., A.B., H.A., A.M.A.

Data Curation: F.A., A.B. Writing - Original Draft: A.B.

Writing - review & editing: F.A., A.B., F.D.v.H. Visualization: F.A., A.B.

Supervision: R.A.B., M.B., F.D.v.H. Project Administration: F.A., F.D.v.H.

Funding Acquisition: A.B., R.A.B., M.B., F.D.v.H.

## Abbreviations

MP: Membrane Proteins
MS: Mass Spectrometry
LC-MS/MS: Liquid Chromatography-Tandem Mass Spectrometry
DDM: N-Dodecyl-β-D-Maltoside
SDC: Sodium Deoxycholate
SP3: Single-Pot Solid-Phase-Enhanced Sample Preparation
SP4: Solvent Precipitation Single-Pot Solid-Phase-Enhanced Sample Preparation
FASP: Filter-Aided Sample Preparation
S-Trap: Suspension Trapping
TEAB: Triethylammonium Bicarbonate
MSP: Membrane Scaffold Protein
spNW30: 30 nm Diameter SpyTag/SpyCatcher Circularized Nanodisc
SMA: Styrene–Maleic Acid Copolymer
SMALP: Styrene–Maleic Acid Copolymer Lipoparticle
SMA2000: 2:1 Styrene-to-Maleic Acid Copolymer
mLiver: Mouse Liver
MASLD: Metabolic Dysfunction–Associated Steatotic Liver Disease
Ni-NTA: Nickel Nitrilotriacetic Acid
LFQ: Label Free Quantification
GRAVY: Grand Average of Hydropathy
FC: Fold Change
LFD: Low-Fat Diet
HFD: High-Fat Diet
MTBE: Methyl Tert-Butyl Ether
MWCO: Molecular Weight Cut-Off
IAA: Iodoacetamide
psi: Pounds Per Square Inch
NSI: nanoelectrospray ionization
FDR: False Discovery Rate
PSM: Peptide Spectrum Match
DDA: Data Dependent Acquisition
TPs: Total Proteins
SPs: Soluble Proteins
MAPs: Membrane Associated Proteins
IMPs: Integral Membrane Proteins
pIMPs: Plasma Membrane Integral Membrane Protein
oIMPs: Organellar Integral Membrane Protein
GO: Gene Ontology
PCA: Principal Component Analysis
TM: Transmembrane Domain
SLC: Solute Carrier

